# Spatial engineering of posterior signaling centers in cerebral organoids via controlled morphogen exposure within hydrogels

**DOI:** 10.64898/2026.06.02.729607

**Authors:** Harrison C. Jeong, Hajime Ozaki, Yuan-Chen Tsai, Caihao Nie, Skylar R. Foust, Kaori Shiraiwa, Ziye Zhuang, Doyle Miller, Matthew Jung Min Noh, Jiya Kinjal Dalal, Aya Galal Salem, Christine Hien Vu, Ali Mohraz, Momoko Watanabe, Herdeline Ann M. Ardoña

## Abstract

Cerebral cortex organoids are powerful *in vitro* models that recapitulate key features of human development. However, conventional methods produce cortical organoids with spontaneous, spatially disorganized cortical regions due to limited control over morphogen distribution within local environments. Here, we present a spatially engineered hydrogel platform that drives localized formation of posterior signaling centers in cortical organoids through controlled, localized exposure to morphogens. Using a combination of bulk photopolymerization, thermal crosslinking, and digital light processing (DLP) approaches, we fabricated hydrogels with stiffness-controlled layers that preferentially deliver morphogens to one side of the organoid, selectively inducing the expression of posterior markers on the exposed face. We further validated this platform by delivering fluorescently tagged dextran, used as molecular weight-matched model morphogens, to visualize spatiotemporal delivery dynamics at the organoid interface. As a proof of principle, we also demonstrated that DLP fabrication enables the printing of dual morphogen hubs, serving as a model for establishing two opposing gradients within a single organoid. Together, this hydrogel platform enables systematic spatial patterning of cell populations in organoids, more faithfully recapitulating the spatial organization and cellular diversity of native tissues and advancing higher-fidelity models for studying human development and disease.

## Introduction

Three-dimensional (3-D) brain organoids, which are *in vitro* tissue reconstructions that recapitulate key architectural, molecular, and cellular features of *in vivo* neurodevelopment, have emerged as powerful models for studying multicellular biology.^1–3^ However, precisely controlling the spatial organization of distinct cell populations to faithfully mirror *in vivo* topographic organization of the brain remains a significant challenge. In native tissue, this topographic organization is directed by specialized clusters of cells called organizers or signaling centers. These spatially restricted organizers release morphogens that form concentration gradients across the tissue.^4–6^ Surrounding cells interpret these positional cues to adopt distinct fates, assembling into the spatially organized tissues that underlie each organ’s specialized function.^4^ Critically, because these gradients originate from the fixed position of the organizer itself, the topographic placement of the organizer is the fundamental determinant of tissue spatial organization. Hence, recreating the organizer topology *in vitro* is essential for generating an accurate topographic structure in organoid models.

In the cerebral cortex, bone morphogenetic proteins (BMPs) and Wingless/Int-1 molecules (WNTs) are secreted from the roof plate organizer, which develops into the secondary organizer known as the dorsomidline telencephalon (DMT), encompassing the cortical hem, choroid plexus, and choroid plaque.^6^ The DMT drives dorsoposteriorization of the cerebral cortex, giving rise to the neocortex, hippocampus, and choroid plexus.^6^ To simulate the process of dorsoposteriorization, previous 2-D and 3-D culture methods have employed bath application of BMPs and/or WNTs (or WNT agonists) to generate dorsal hippocampal and choroid plexus cells from human pluripotent stem cells (hPSCs).^2, 7–9^ Similarly, our previous study demonstrated that bath application of BMP4 and CHIR-99021 (a canonical WNT agonist) was essential for generating the DMT, which in turn posteriorized the adjacent neocortex within cortical organoids toward visual cortex-like areas.^10^ BMP4 alone is sufficient to induce the DMT,^7^ but the addition of CHIR-99021 has been proven to support DMT formation.

To better control the presentation of morphogens to *in vitro* cortical models, previous studies attempted to pattern neural features with a morphogen gradient using microfluidic devices.^11–14^ Instead of generating a heterogeneous collection of organoids with varying identities,^13^ our previously reported approach generated a single, topographically organized tissue that recapitulates the continuous dorsoventral axis within a unified architecture.^14^ This approach affords region-specificity in cortical organoids, unlike other approaches focused on elongated neural tube tissue ^11, 12^ that covers the entire anteroposterior axis of the CNS. However, these cortical organoids are small and spherical, making it difficult to control (i) the precise location of the organoid within a large chamber in a microfluidic chip; (ii) the sustainability of a morphogen gradient within a large chamber in a chip; and (iii) the scalability of the organoid production since only one organoid was placed in each microfluidic chip to create a fine morphogen gradient.^14^

To overcome these technical challenges, one approach is to design synthetic matrices that will allow for the initial localization of morphogens to induce controlled delivery from engineered matrices and, consequently, elicit spatially defined responses. This approach has been broadly applicable to the local delivery of drugs and other chemical cues across different types of *in vitro* models. For example, spatial control of biochemical cues has been demonstrated across multiple contexts, including differential endothelial cell attachment via gradient presentation of immobilized epitopes on hydrogel surfaces,^15^ improved axonal extension across injured environments through local paclitaxel release from electrospun polymer matrices,^16^ and induction of neural organoid regionalization through a single morphogen-gradient generator device.^17^ These examples highlight the tunability offered by synthetic matrices in controlling chemical cue delivery based on the physical properties of such artificial environments. However, simultaneously maintaining viable culture conditions for cortical organoids while achieving spatially resolved exposure to multiple morphogens in confined matrices remains a significant challenge, particularly when scalable manufacturing and high-throughput organoid production are considered.

Here, we present a facile and scalable polymeric hydrogel-based approach for controlled regionalization of cerebral cortex organoids. To dictate the localization and timing of morphogen exposure of organoids, we developed multi-layered hydrogel bioscaffolds with three compartments (from top to bottom): an organoid layer, an intermediate layer, and a morphogen hub–with respective geometries and stiffnesses that are spatially controlled using light-based patterning. These polymeric hydrogel scaffolds are designed to be compatible with standard cortical organoid culture conditions and digital light processing (DLP) 3-D bioprinting, enabling reproducible fabrication without the need for complex microfabrication. Stiffness differences across compartments were used to tune morphogen diffusion kinetics throughout the organoid encapsulation period. Posterior signaling center formation was then characterized and compared between organoids exposed to morphogens uniformly versus directionally within the hydrogel environment. Overall, this work presents a spatially engineered approach in presenting morphogens to promote anatomically and physiologically relevant cortical organoids, ultimately supporting improved cellular diversity, circuit maturation, and a better understanding of neurodevelopment and disease.

## Results

### Establishing cortical organoid-compatible composite hydrogels

First, we screened for hydrogelators and crosslinking conditions that enable a sustained culture of Day 15 cortical organoids while exposing them to posterior morphogens (BMP4 and CHIR-99021) over a 6-day period **(Figure 1A–B)**. We previously showed that this 6-day period was an optimal window to induce anterior and posterior signaling centers that eventually self-polarize the adjacent tissue into antero-posterior areas in cortical organoids.^10^ Our selected set of polymers/polymer blends capable of stimuli-induced gelation (e.g., temperature or light) focuses on commercially available materials to facilitate scaling of the fabrication approach presented here. These included commercially available: Mebiol® (a thermoresponsive poly(*N*- isopropylacrylamide)-poly(ethylene glycol) copolymer, PNIPAAm-PEG), HAMA (methacrylated hyaluronic acid), PEGDA (diacrylated PEG), and GelMA (methacrylated gelatin). Blends of these components, such as Mebiol:HAMA (M:H) and Mebiol:PEGDA (M:P), were also included in the screening panel. Because radical species generated during photopolymerization can limit the viability of organoids in DLP-based bioprinting, we optimized the UV irradiation duration to preserve organoid viability comparable to bath conditions (control) while still achieving self-supporting gel scaffold formation. To quantitatively assess organoid viability during material screening, organoid morphology and cleaved caspase-3 (cl-CASP3, standard biomarker for apoptosis) expression, were measured to confirm that the light-induced hydrogelation and bioprinting conditions did not significantly induce cell death **(Figure 1C–E)**. Based on these parameters, a direct 1-minute exposure to 405 nm (10 mW/cm^2^, wavelength used for photopolymerizations here) resulted in organoids with no significant deviation from the organoid morphology or cl-CASP3 levels of the control/untreated condition, suggesting this short UV exposure did not lead to excessive cell death in organoids. Among the material compositions screened, the formulations with Mebiol as a major component consistently showed an intact epithelial layer of the organoids. Interestingly, mixing PEGDA as a photopolymerizable component with the thermoresponsive Mebiol did not help with maintaining the desired organoid morphology features. Even at 9:1 (v/v) Mebiol:PEGDA (M:P) ratio, this condition showed about a 2-fold increase in cl-CASP3 apoptotic marker levels as compared to the control and UV-irradiated organoids for 1 minute **(Figure 1E)**. We also note that the stability of interlayer adhesion is a critical factor for the constructs. Lowering the PEGDA component below 10% leads to higher probability of layer dissociation throughout the 6-day culture. Considering all the criteria for screening imposed here, and the need to have photocrosslinkable groups in the ink formulation for DLP-based printing down the line, Mebiol:HAMA 9:1 (v/v) was deemed the most suitable hydrogel encapsulation of cortical organoids.

**Figure 1.**
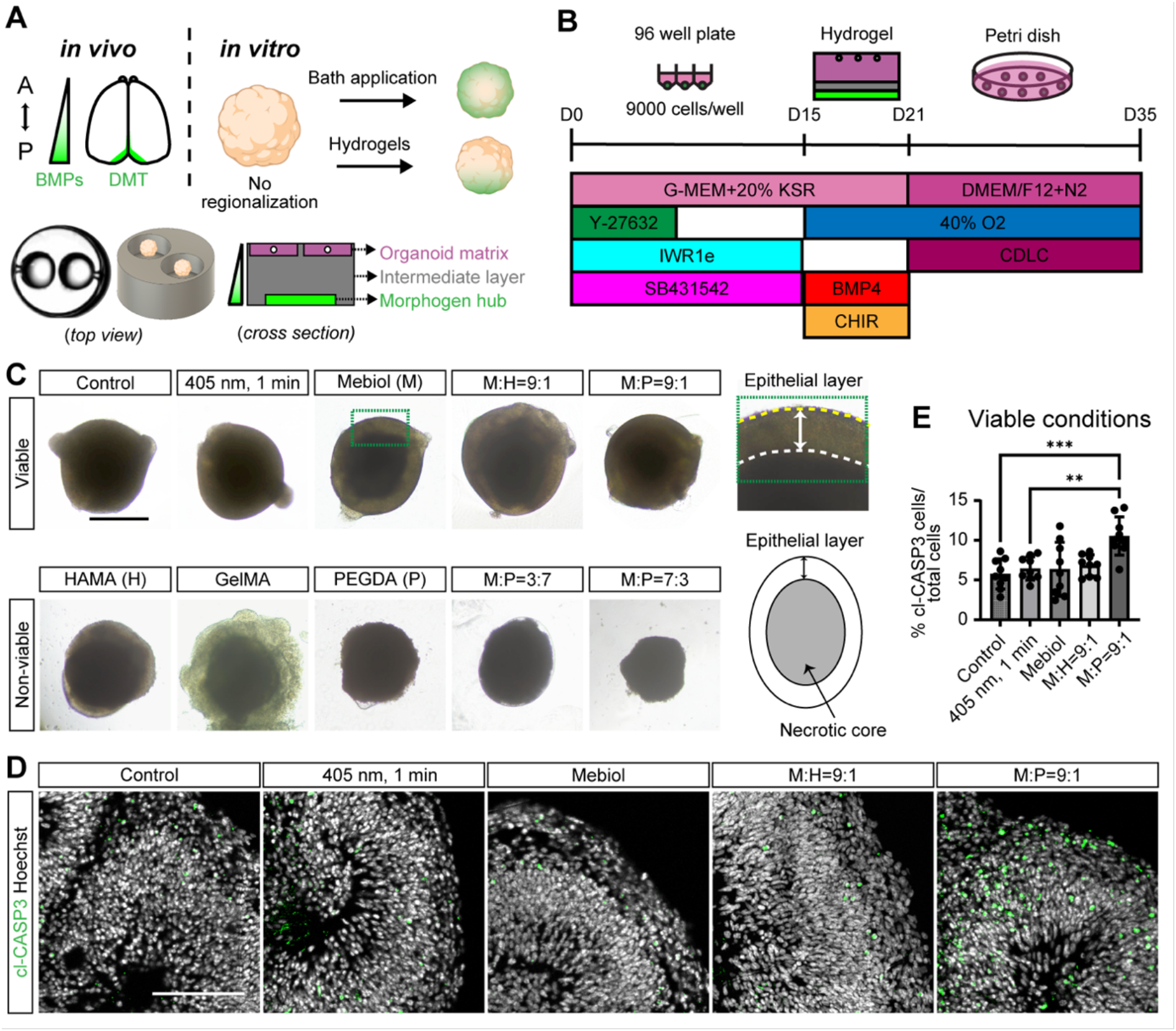
Experimental schematic and hydrogel compatibility screening for cortical organoid culture. (A) This work presents a hydrogel-based *in vitro* model of the *in vivo* cortical regionalization. (B) Timeline of organoid culture conditions utilized for this study. (C) Representative images showing healthy versus unhealthy organoid morphology at Day 21, after 6 days of encapsulation under different hydrogel conditions. Control: organoid in standard media. 405 nm light: organoid exposed to light for 1 min in standard media. All photocrosslinkable hydrogels were polymerized with 405 nm light for 1 min. M:H= Mebiol and HAMA blend; M:P= Mebiol and PEGDA blend. Scale bar: 500 µm. (D) Immunostaining images showing a cell death marker, cl-CASP3, with nuclear staining, Hoechst. Scale bar: 100 µm. (E) Quantification of cl- CASP3 positive cells in cortical organoids embedded in different viable hydrogels at Week 5 from 2 independent hPSC (H9 and XF) lines with at least 4 experimental replicates (N= 9 organoids per condition). Data are represented as mean ± SD. The p-values were calculated using one-way ANOVA with Tukey’s post-hoc test. **p<0.01; ***p<0.001.

### Visualization and modeling of morphogen diffusion

Based on the pre-screened polymer compositions from Figure 1, we studied the macromolecular diffusion profiles across different hydrogel platform designs over a 6-day period, a critical period for inducing posterior signaling centers.^10^ PEGDA was used as the photocrosslinkable morphogen hub component, while the M:H (9:1) blend was selected as the photocrosslinkable and thermoresponsive organoid matrix layer based on the viability screening. M:P was selected for the intermediate layer to contain components similar to those in the top and bottom layers, and to include a photocrosslinkable component to help stabilize the interface between the layers. To visualize diffusion patterns by confocal fluorescence microscopy, FITC-dextran (40 kDa), a fluorescently tagged molecule with a similar molecular weight to dimeric BMP4, was used as a model morphogen **(Figure 2** **and S1–S5)**. CHIR-99021 is considerably smaller than BMP4 and is therefore expected to diffuse more readily. We focused our analysis on BMP4, as it is sufficient on its own to induce DMT formation.^7^ The three hydrogel models studied are as follows: (#1) a uniform M:H hydrogel encapsulating 400 or 0 ng/mL of FITC-dextran, as a positive or negative control, respectively **(Figure 2A)**; (#2) a two-layered platform with a 1 mm PEGDA morphogen hub layer (4000 ng/mL FITC-dextran) beneath a M:H (9:1) top organoid matrix layer **(Figure 2B)**; and (#3) a three-layered platform similar to Model #2, but with an additional 1 mm M:P (1:9) intermediate layer situated between the bottom morphogen hub layer and the top organoid matrix layer **(Figure 2C)**. Under all three hydrogel model conditions, no organoids were added in the topmost layer throughout the fluorescence modeling studies. The 2- D fluorescence heatmaps across the coronal plane of Models #1 to #3 demonstrate the impact of each hydrogel configuration in slowing down the transport of FITC-dextran from the bottom hub to the topmost layer. The positive control (#1) exhibited a significant reduction in fluorescence relative to the morphogen hub conditions (#2-3) at initial timepoints, suggesting that FITC-dextran readily diffuses from the hydrogel construct into the surrounding media. At Day 6, the 2-D heatmaps and 3-D fluorescence reconstruction data showed clear differences in FITC-dextran distribution between Models #2 vs. #3 across the z-axis (*i.e.,* from dotted yellow to green lines) once M:P was added as a resistive intermediate layer **(Figure 2B–C)**. To demonstrate release of the morphogen itself rather than of a dextran surrogate, BMP4 released into the culture medium from hydrogels prepared as Models #1, #2, and #3 was quantified by ELISA **(Figure S6)**. The protein concentrations measured in the culture medium atop the hydrogels are consistent with the trends observed in the fluorescence visualization experiments: BMP4 accumulation was lowest for Model #3, which incorporates an intermediate layer, relative to Model #2, which lacks the intermediate layer. These results support our claim that the intermediate layer restricts morphogen transport and thereby confers spatial control over the amount of morphogen that reaches the top hydrogel region.

**Figure 2.**
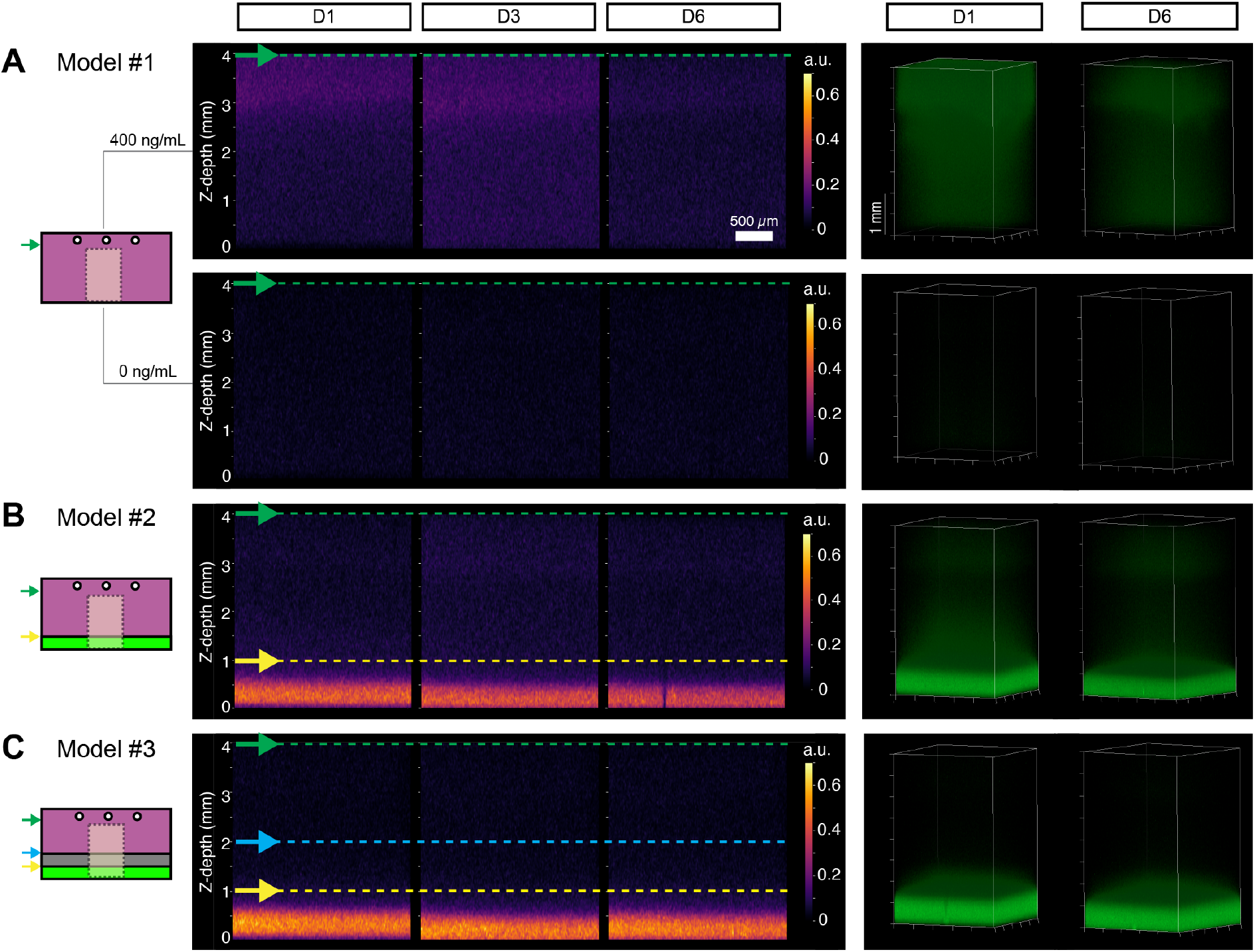
Visualization of varied morphogen diffusion profiles across hydrogel models. Schematics on the left illustrate the three model hydrogel conditions used for morphogen diffusion studies using fluorescence monitoring. FITC-dextran (40kDa) was used as the fluorescent macromolecule to model morphogen diffusion across hydrogel conditions. The region within the dashed line represents the imaging area for each model schematic. (A) Model #1- M:H (9:1) composite gel (pink) uniformly encapsulating 400 (positive control) or 0 (negative control) ng/mL FITC-dextran; (B) Model #2- two-layered hydrogel with localized FITC-dextran in PEGDA layer (green, 4000 ng/mL) in the bottom and organoid-compatible M:H layer at the top (pink); (C) Model #3- 3-layered hydrogel comprised of Model #2 layers with a middle M:P (1:9) layer (grey). Middle: Representative FITC-dextran fluorescence heatmaps across the central coronal plane of the bulk hydrogel construct, taken at different timepoints throughout the 6-day period. Scale bar: 500 μm. Right: 3-dimensional reconstruction of FITC-dextran fluorescence levels. Scale bar: 1 mm.

To further rationalize the ability of the additional layers to effectively slow down the transport of a 40 kDa macromolecule across the hydrogel system, we characterized the mechanical properties of each layer through parallel plate oscillatory rheological measurements **(Figure 3** **and S7)**. It is expected that the diffusivity of the macromolecule will be lower in stiffer hydrogels with higher polymer network and crosslink densities. These measurements also allow us to compare the stiffness of the relevant layers with the physiological stiffness of brain regions.^18, 19^ Frequency sweeps within the linear viscoelastic region for all tested materials were performed to obtain the storage and loss moduli (G′ and G″) values for each hydrogel. The frequency sweep data show that at 1 Hz, PEGDA is the stiffest material (G′ ∼61 kPa), followed by M:P 1:9 (G′ ∼19 kPa), and M:H 9:1 is the softest (G′ ∼0.2 kPa) (**Figure 3A**). We used 1 Hz data to represent material behavior that falls within the tested frequency range (0.1 to ∼30 Hz), while recognizing that the approximations for simulations discussed below may slightly change at other frequencies due to the increasing M:H moduli under these conditions. Comparing the three hydrogels, we confirmed that the measured stiffness of M:H hydrogel used here is most relevant for our organoid experiments because it is similar to that of the central nervous system tissue.^18, 20–24^ Based on the information about the relative stiffness of the layers that constitute each hydrogel scenario for the simulations, an analytical and numerical model of morphogen diffusion was developed whereby all variables (η= full length of mass transfer zone; θ=morphogen concentration; *τ*= diffusion timescale; b= height of the morphogen hub) are non-dimensionalized. Further details on the definition of these variables, as well as the solutions used for both analytical and numerical models for all simulated diffusion scenarios can be found in the Supplementary Information. The relative diffusivities across the morphogen hub layer, intermediate layer, and organoid matrix layer were then approximated as 0.003, 0.011, and 1, respectively, based on relative storage moduli values measured in Figure 3A. Using the analytical and numerical models setup for our hydrogel system, θ fully equilibrates across the system after approximately one diffusion timescale and beyond (*τ* ≥ 1) for the control hydrogel Scenario #1 (**Figure 3B**). The addition of another layer with ten times less diffusivity (Scenario #2) shows a slowdown of transport as compared to the theoretical hydrogel with uniform diffusivity in Scenario #1 (**Figure 3C**). The diffusion progressively slows down upon the addition of a middle resistive layer in Scenarios #3 and #4, recapitulating the addition of the M:P intermediate layer in experimental hydrogel #3 but with varied thicknesses (**Figure 3D–E**). At η=0.8, where the morphogen diffusion front is estimated to start interfacing with organoids, the θ in Scenario #2 becomes about a tenth of Scenario #1 after a tenth of a diffusion cycle (τ∼0.1) (**Figure 3F**). The progressive slowdown of morphogen transport becomes more apparent when stiffness differences between layers and an intermediate layer are introduced. It is worth noting that in Scenario #4, when majority of the construct volume is comprised of the intermediate layer, θ values approach 0 at all τ. Experimentally, the decreasing trend in FITC-dextran transport between #2 and #3 (**Figure 2B–C**) is consistent with the simulations in Figure 3 but is more pronounced in the experiment. Collectively, these results help rationalize the experimentally observed slowdown in morphogen transport from the hub layer to the organoid layer due to stiffness differences between hydrogel layers. Scenario #2 to #4 simulations also demonstrate the importance of optimizing the volume fraction of each distinct layer to achieve a reasonable concentration of the morphogen at a specific time and to not completely impede transport towards the organoid interface. The simulated scenarios further approximate the fraction of morphogen reaching the organoid interface after one diffusion cycle under each layering condition.

**Figure 3.**
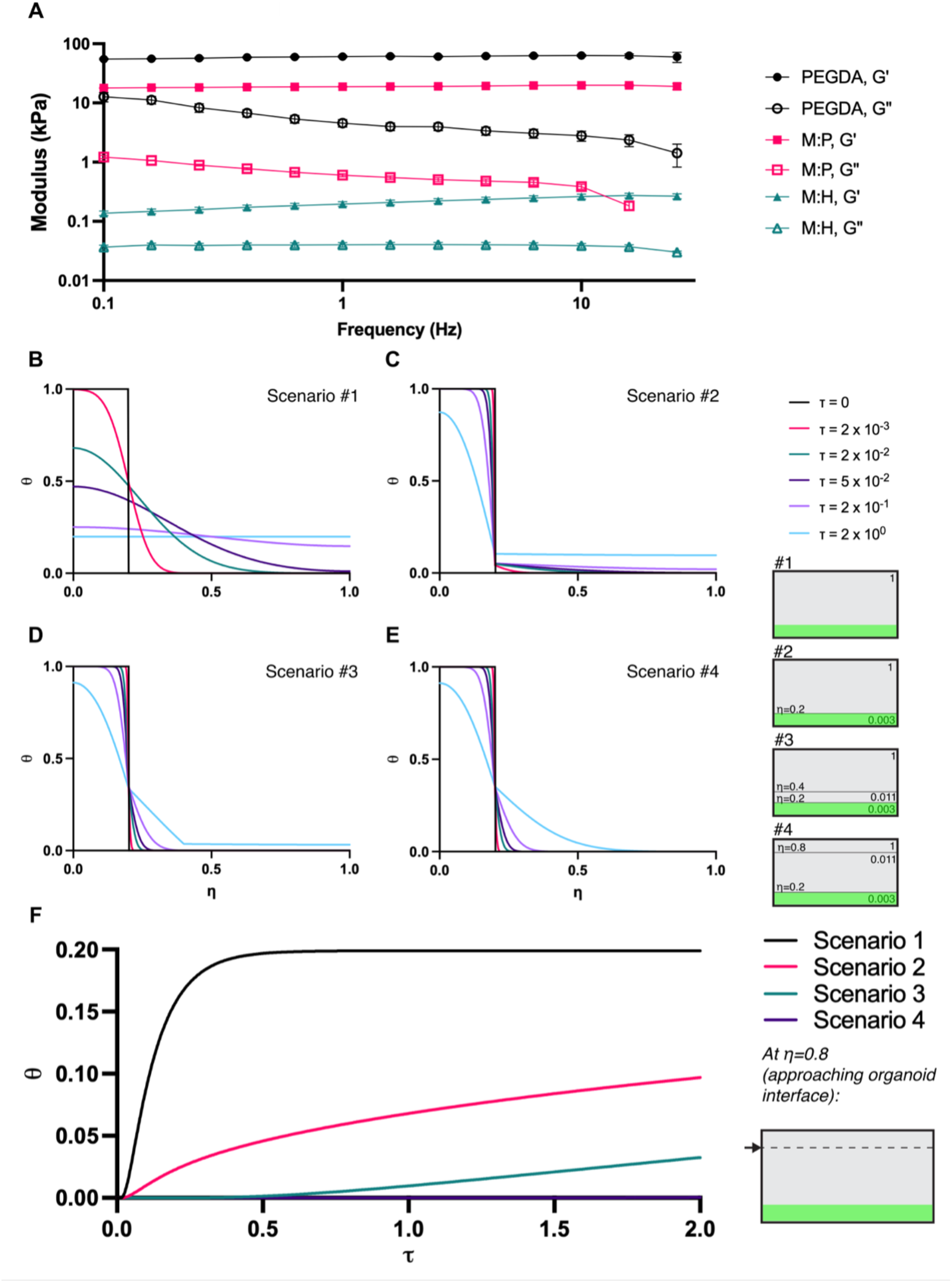
Simulating morphogen diffusion across theoretical layers of varied stiffnesses. (A) Frequency sweep measurements at 0.1% strain at 37°C for the hydrogels used in fluorescence modeling experiments in Figure 2 (N= 3; error bars represent standard error of the mean, s.e.m.). (B-E) Simulating morphogen diffusion across hydrogels under different scenarios (#1–#4); normalized morphogen concentration (θ) vs. normalized diffusion lengths (η) at different diffusion timescales (τ). Schematics on the right represent the relative diffusivities between the pertinent hydrogel layers (top, middle, bottom; 1, 0.011, 0.003, respectively). The assigned relative diffusivities mimic the stiffness differential between the M:H, M:P, and PEGDA layers used in the experiments. In all scenarios, morphogen is initially loaded in the region η = 0–0.2 (green shaded region). (F) θ vs. τ curves extracted at η = 0.8 to simulate morphogen concentration at the organoid interface across all scenarios.

### Spatially directed induction of posterior signaling centers within layered hydrogels

To confirm if posterior signaling centers can be induced in our hydrogel models, cortical organoids were first embedded in Mebiol hydrogels (similar to Model #1 setup) uniformly encapsulating 0–400 ng/mL of BMP4 and 0–240 µM of CHIR-99021. Specific concentrations of CHIR-99021, paired with BMP4 concentrations, are noted in Materials and Methods. Our data showed that 50, 100, 200, and 400 ng/mL BMP4 encapsulated in Mebiol hydrogels effectively induced the posterior signaling center DMT, marked by OTX2 and TTR. The negative control, the Mebiol hydrogel with no BMP4 (0 ng/mL), did not significantly induce DMT (**Figure S8**). This result is consistent with the bath application dose ranges validated in our previous publications^10, 14^ and suggests that morphogen responsiveness of cortical organoids is largely preserved within the Mebiol polymer network. Next, cortical organoids were exposed to the posterior morphogen BMP4 using the established hydrogel multi-layered platforms. For these experiments, it is important that the organoid location and orientation are fixed within the hydrogels throughout the 6-day culture. All organoids that were processed for data analysis satisfied this criterion. In Model #1, positive control hydrogels, organoid-compatible M:H hydrogels encapsulating 400 ng/mL of BMP4 upregulated posterior signaling center DMT markers (OTX2, TTR, and MSX1) and downregulated a cortical marker PAX6, compared to those without growth factors, as confirmed by immunochemical and RT-qPCR assays **(Figure 4A–C** **and S9)**. Furthermore, we typically recognized more than one signaling center region in Model #1 positive control hydrogels **(Figure 4E)**. In Model #2 hydrogels, when cortical organoids were incorporated into the upper M:H hydrogels and BMP4 (400 ng/mL or 4000 ng/mL) was confined to the lower PEGDA morphogen hub, no significant increase in posterior signaling center markers was observed **(Figures 4A–C)**. This may be attributed to the rapid diffusion of BMP4 from the multilayered structures into the surrounding media, which was replaced daily. In Model #3, both immunofluorescence imaging and marker quantification data clearly verified the formation of one posterior signaling center region that can be induced when a higher concentration of BMP4 (4000 ng/mL) was localized in the PEGDA morphogen hub, indicating a spatially directed induction of a posterior signaling center within one cortical organoid **(Figures 4A–C)**. All other subsequent experiments comparing morphogen introduction via Model #2 and #3 used 4000 ng/mL in the morphogen hub layer. This observed phenomenon can be attributed to the presence of an intermediate M:P layer, which contributes to the slower diffusion rate of BMP4 throughout the hydrogel construct as shown in Figure 2 and simulated in Figure 3.

**Figure 4.**
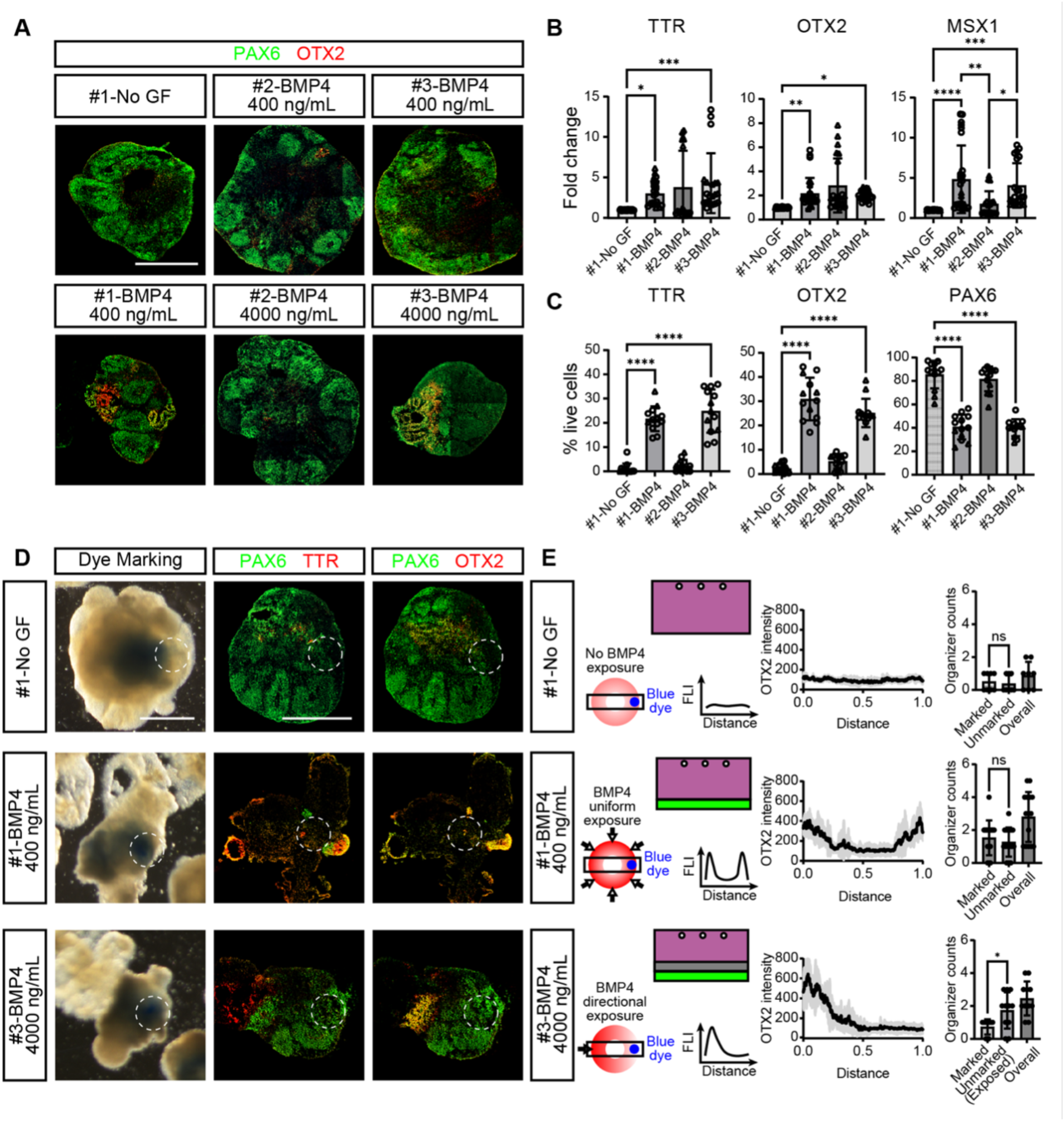
Induction of posterior signaling centers within cortical organoids embedded in Models #1–#3. (A) Immunochemical analyses showing cortical (PAX6) and posterior signaling center (OTX2) markers. Scale bar: 500 μm. (B) Quantification of posterior marker induction by RT-qPCR (from H9 hESC and hIPS2 hiPSC; N= 15–18 samples per condition, derived from 4–5 pooled organoids per sample, 6–7 independent batches). BMP-high: BMP4 4000ng/mL for H9 and 2000ng/mL for hIPS2. (C) % positive cells out of total live cells quantified from the immunofluorescence images in A (from H9 hESC and hIPS2 hiPSC; N= 17 organoids per condition, 6–8 independent batches). (D) The upward (unexposed) side of the organoids was marked by Brilliant Blue G dye (white dotted circles). Scale bar: 500 μm. (E) Quantification of posterior signaling center induction on the marked and unmarked (exposed) sides of individual organoids. OTX2 fluorescent intensity was measured along the organoid diameter and plotted over normalized distance (0–1). Posterior signaling center counts based on TTR and OTX2 expression were quantified per organoid section, revealing preferential induction on the BMP4-exposed side in Model #3 (N= 8–15 organoids per condition, 3 independent batches). Data are presented as mean ± SD. The p-values were calculated using one-way ANOVA with Tukey’s post hoc test. *p<0.05; **p<0.01; ***p<0.001; ****p<0.0001.

To further verify whether the exposed side of the organoid differentiated into a posterior signaling center, we labeled the upward-facing side (unexposed) surface of the organoid with Brilliant Blue G (BBG) dye upon organoid retrieval from the hydrogels after 6 days of exposure at Day 21 **(Figures 4D–E** **and S10)**. We kept each organoid in a well of the 96-well plate to track the marked unexposed side based on the morphology, even if the dye was faint at Day 35 when we characterized the organoids. We mounted the unexposed side on top, so we can identify the marked and unmarked sides of the organoid. For cortical organoids with no GF (Model #1, negative control, no BMP4), we measured a few spontaneous inductions of DMT, but the location of the induction was random **(Figure 4D–E)**. For cortical organoids with localized BMP4 in the PEGDA morphogen hub (Model #3, 4000 ng/mL BMP4), we detected the generation of DMT at high coincidence with the exposed side, showing characteristic papillary tissue formation for one of the DMT structures, the choroid plexus^7^ **(Figure 4E)**. Comparing the three hydrogel models studied here, only Model #3 with localized BMP4 showed spatially directed induction of DMT.

We sought to demonstrate that the specific placement of organoids with respect to the morphogen source location can be further controlled via DLP-based printing of microwells within the organoid matrix layer. The utility of DLP as a fabrication technique here also offers a reliable way to scale up the production of such hydrogel constructs in a matter of minutes. As such, Models #4 and #5 **(Figure 5A)** were developed as hydrogel conditions to illustrate the DLP-printed analogs of Model #3, where the bottom and top layers are localized as smaller discs within the center of the construct. We performed the FITC-dextran diffusion experiment for both DLP-printed models using Model #4, since the only physical difference between the two is the addition of a second microwell in the upper region of the printed gel. In Model #4, the FITC-dextran diffusion profile is similar to Model #3, whereby a slowdown of FITC-dextran transport is apparent in comparison with Model #2 **(Figure 5B** **and S11)**. Upon seeding organoids onto these hydrogel configurations, the same trends for localized markers were observed for organoids in #4 and #5 as compared to the negative control condition in Model #1 **(Figure 5C–D** **and S12)**. The posterior signaling center DMT markers TTR and OTX2 were both upregulated when organoids were subjected to Model #4 or #5 hydrogels, as compared to the Model #1 negative control. Similarly, the cortical marker PAX6 remains elevated in Model #1 as compared to the posteriorized organoids in Model #4 and #5.

**Figure 5.**
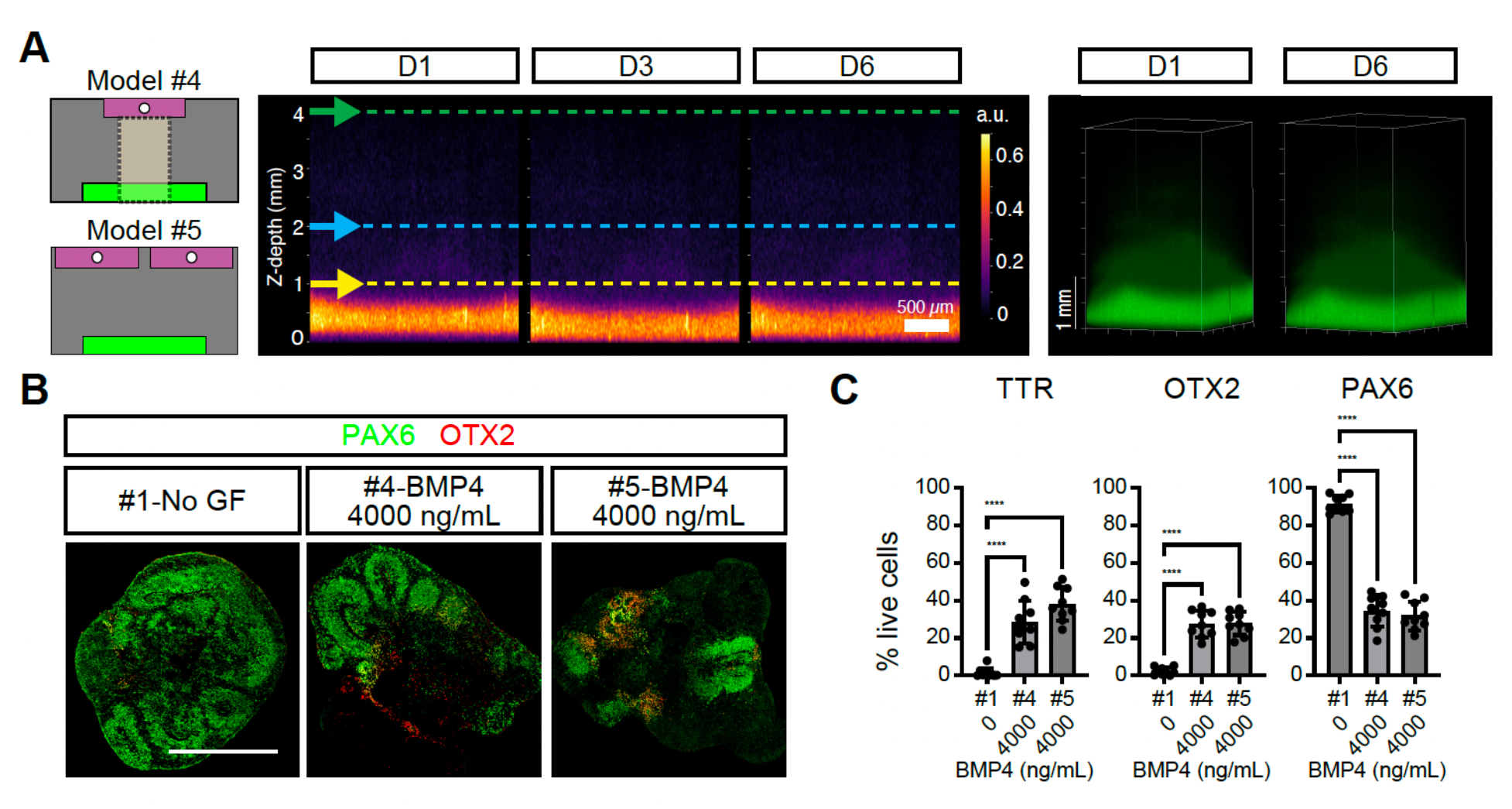
Induction of posterior signaling centers within cortical organoids embedded in DLP-printed, multi-compartment hydrogels. (A) Visualization of FITC-dextran diffusion across DLP-printed hydrogels (Models #4 and #5). Dashed region represents the imaged area in Model #4. Scale bar: 500 μm. (B) Immunochemical analyses showing cortical (PAX6) and posterior signaling center (OTX2) markers. Scale bar: 500 μm. (C) Quantification of TTR-, OTX2- and PAX6-positive cells from immunostaining analyses, expressed as the percentage of marker-positive cells among total Hoechst-positive cells (N = 9 organoids per condition, 3 independent batches). Data are presented as mean ± SD. The p-values were calculated using one-way ANOVA with Tukey’s post hoc test. ****p<0.0001.

### Tuning morphogen diffusion profiles via DLP-endowed hydrogel characteristics

Another hydrogel property that was investigated to potentially tune the morphogen diffusion profile is the establishment of stiffness gradients. Here, we imposed a light intensity gradient as the xy-planes are being DLP-printed for hydrogel constructs with similar geometry to Model #5 **(Figure 6A** **and S13)**. A grayscale image of a circle with gray values ranging from 100% to 50% white for the photomask (across the x-direction) was used, and a printhead speed of 0.01 mm/s was maintained throughout the entire body. Rheological measurements performed on M:P hydrogels generated using uniform DLP photomasks with different gray values (100%, 75%, and 60%) confirm the impact of the graded mask on the stiffness of the printed constructs **(Figure S14)**. For reference, the M:P intermediate layers in Models #3, #4, and #5 hydrogels were exposed to 60% light intensity. The FITC-dextran concentration was kept at 4000 ng/mL for the morphogen hub layer. The Day 0 transverse plane images represent the starting conditions where morphogens are confined in the morphogen hub layer. After 1 day of incubation, we observed that the regions of higher intensity at Z=380 µm transverse plane is not the same as the regions of higher intensity at a higher transverse plane (e.g., Z=980 µm). If this was due to heterogeneity in morphogen loading as seen in some hydrogels with homogeneous stiffness **(Figure S11)**, the regions of high fluorescence intensity should be consistent throughout the Z-direction. In the case of hydrogels printed with a gradient photomask, the representative Day 1 images in Figure 6A show that the higher fluorescence intensity regions in Z=380 µm plane is not the same as that in Z=980 µm plane. At lower Z-planes, the higher intensity regions seem to represent the stiffer, more polymerized regions where morphogens diffuse less axially. At higher Z-planes, the higher intensity regions can be attributed to the less stiff, less polymerized regions where morphogens can diffuse more axially. The same pattern of variation in regions of higher intensity persists at Day 6 of incubation.

**Figure 6.**
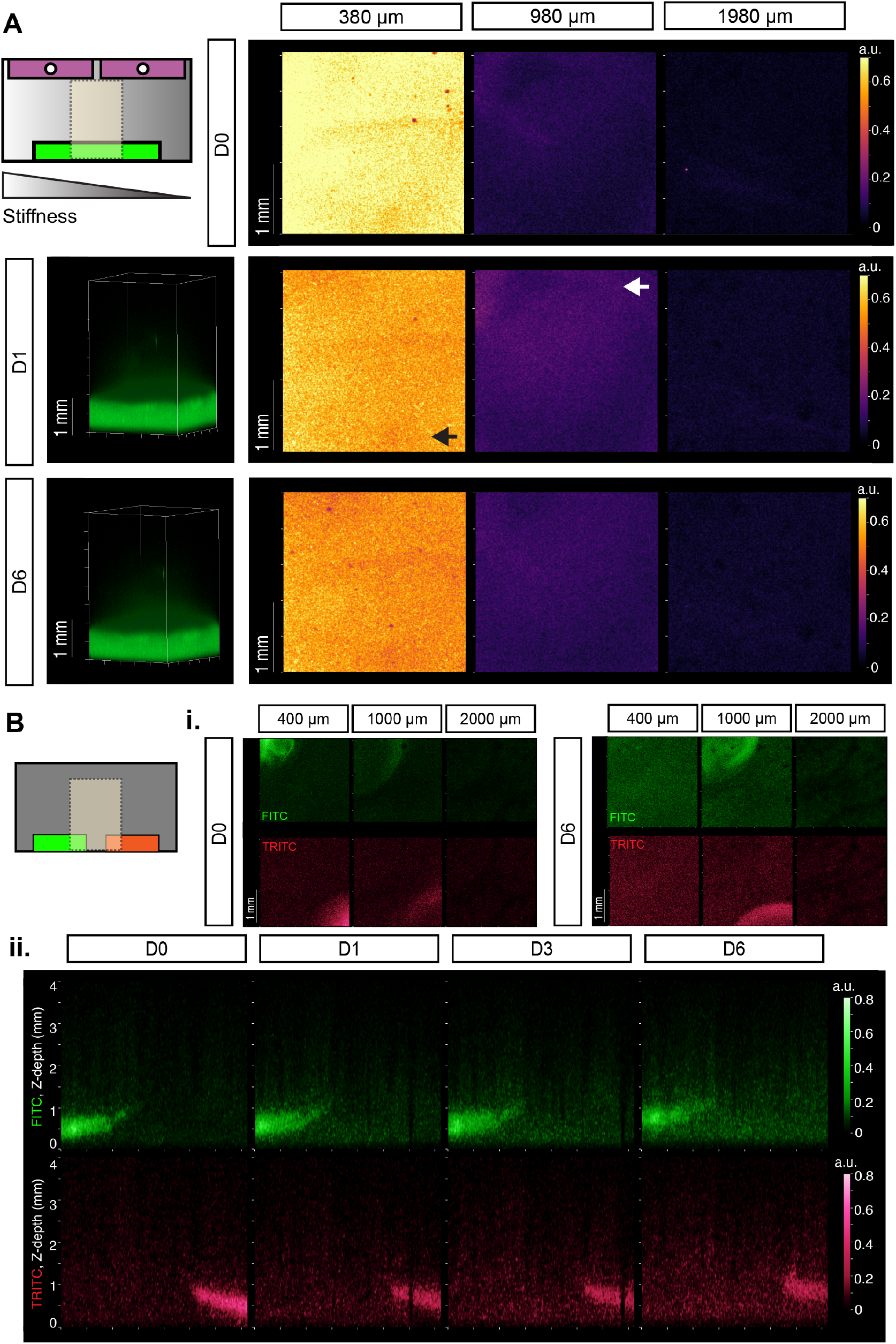
Tuning morphogen diffusion profiles and localization using DLP-dictated hydrogels properties. (A) Representative diffusion heatmaps across different transverse planes (Z= 380, 980, 1980 µm) for a DLP-printed hydrogel construct with lateral stiffness gradient. The morphogen hub layer is loaded with 4000 ng/mL FITC-dextran. Arrows in the Day 1 transverse-plane fluorescence maps highlight regions of higher intensity. (B) Representative fluorescence images (i) across different transverse planes (Z= 400, 1000, 2000 µm) at Day 0 and Day 6; and (ii) across a central coronal plane for a hydrogel construct DLP-printed to localize two morphogen hubs, which are modeled by FITC- and TRITC-dextran. For both hydrogel construct schematics in (A) and (B), the dashed region represents the imaged area.

As another demonstration of DLP-endowable hydrogel characteristics that can serve as a precursor to future work on controlled multi-morphogen exposure, we designed a DLP-printable hydrogel construct for a two-morphogen system. To model the inclusion of antagonistic morphogens within our hydrogel platform, we monitored the diffusion of 40 kDa FITC-dextran and TRITC-dextran (similar to the size of BMP4 and an anterior-inducing morphogen, FGF8 ^2, 10, 25^ throughout our model system over a 6-day experimental period **(Figure 6B** **and S15–20)**. We fabricated this model by first printing a cylindrical M:P base with two open regions. Each open region was filled with FITC- or TRITC-dextran in a PEGDA solution. The solutions were then crosslinked via direct exposure from a 405 nm lamp. Following the polymerization of these materials, the rest of the M:P gel was printed on top of the preliminary base. Figure 6Bi–ii demonstrates the resulting localization of each model morphogen within DLP-printed hydrogels with light-sculpted microwells that house the FITC- and TRITC-dextran separately. Similar to Models #4 and #5, morphogen diffusion was apparent over the 6-day period, and the presence of morphogen within the concentrated hubs decreased with time **(Figure 6Bi)**.

## Discussion

A central challenge in organoid biology is the absence of spatially controlled morphogen delivery and spatially directed differentiation. Conventional approaches relying on bath application of signaling molecules to the culture medium sacrifice spatial precision, as uniform morphogen dispersal promotes spontaneous differentiation that lacks the topographic organization characteristic of native tissue. This limitation applies to current approaches for cerebral cortex organoid production, whereby posterior signaling center induction in most existing protocols is random and uncontrolled. In this work, we presented an approach to control posterior signaling center induction in cortical organoids using an engineered hydrogel platform compatible with light-based fabrication techniques such as DLP. We note that the full transcriptional and functional characterization of posterior signaling centers, including single-cell RNA-sequencing, spatial transcriptomics, and conditioned medium assays, was previously reported in the same cell line and culture system;^10^ we therefore identify them here using representative markers. The polymer/polymer blend compositions for the hydrogels used here were selected based on organoid compatibility and the ability to crosslink in response to external stimuli, such as light (*i.e*., for photopolymerizability and DLP compatibility) or heat (*i.e*., providing a practical advantage for organoid encapsulation at room temperature before incubation at 37°C). Results from the cl- CASP3-based organoid viability assessments **(Figure 1)** confirmed that the material selection and polymerization conditions used did not adversely affect organoid health. The photocrosslinkable polymeric components were also retained to stabilize adhesion at the interface between each layer. The overall processing conditions for each layer were optimized to establish sufficient stiffness differential within the hydrogel construct, while allowing for the organoid matrix layer (M:H) to match the mechanical properties critical for organoid viability and function.^26^ These fabrication considerations led to the hydrogel-based platform described here aimed at addressing limitations in controllable morphogen exposure localization and timing for cortical organoid regionalization.

Fluorescence-based tracking of dextran diffusion revealed the capacity of stiffness- and geometry-defined hydrogel layers to regulate morphogen transport **(Figures 2, 5, 6)**. The DLP-printed platform design provides additional spatial control over organoid positioning due to the presence of microwell features **(Figure 4)**. In both manually photopolymerized and DLP-printed hydrogel constructs, the conditions where an intermediate layer was added to slow down morphogen transport are the ones that afforded localized posterior signaling center induction on the morphogen-exposed organoid surface **(Figures 4 and 5)**. Notably, even with the increased intermediate layer thickness in Models #4 and #5 as compared to Model #3, organoids continued to develop localized posterior signaling centers **(Figure 4)**. The experimentally observed diffusion profiles across hydrogel conditions of varied stiffness were further supported by analytical and numerical simulations developed for this system **(Figure 3)**. Together, the collective modeling and simulation results provide a future foundation for further refinement of the platform and for modulating the extent of posterior signaling center induction. The insights from these simulations also offer a general platform for investigating a broader design space for experimentally generating such multi-layered/-compartment hydrogels as a delivery device for controlled exposure of organoids to morphogens.

Finally, the fabrication of hydrogels with stiffness gradients and two morphogen hubs represents an important step towards creating cortical organoids with greater physiological complexity. During development, organs rely on multiple spatially distinct organizers that secrete different signaling molecules at varying concentrations and in a coordinated manner. In the cerebral cortex, for example, the anterior organizer (anterior neural ridge) secretes FGFs, while the posterior organizer (DMT) secretes BMPs and WNTs—pathways known to interact antagonistically.^27^ While the organoid assays shown here were limited to the one-morphogen system (BMP4), the demonstration with the two-morphogen hydrogels specifically illustrates the feasibility of incorporating multiple morphogen hubs within a single platform to study the effects of controlled, spatially directed delivery of antagonistic morphogens. Ultimately, the signaling landscape governing cortical development is considerably more complex, involving a larger repertoire of molecular cues and cellular responses. While we only show here the hydrogel fabrication conditions to resolve the transport of two macromolecules with similar molecular weights across the hydrogel constructs, further optimization is required to co-deliver the appropriate concentrations of antagonistic morphogens that will be effective for the concurrent posteriorization/anteriorization of single organoid units. The long-term goal of this work is to develop a scalable, bioprintable hydrogel platform capable of recapitulating a greater degree of this complexity, bringing *in vitro* models closer to the physiological organization of native tissue. Overall, we anticipate that this work will lay the groundwork for more spatially defined differentiation strategies, ultimately supporting greater cellular diversity, improved neural circuit maturation, and a deeper understanding of neurodevelopmental processes and disease.

## Materials and Methods

### Bulk or layer-by-layer assembly of hydrogels for Models #1–#3

For bulk hydrogels (#1-3), the hydrogelators at liquid state were deposited on a well plate on ice. Lithium phenyl-2,4,6- trimethylbenzoylphosphinate (LAP, Advanced Biomatrix, #5269) was used as a photoinitiator for all hydrogels and dissolved in PEGDA solution (0.05 wt%) or SAS-CD (1.7 wt%) for M:H gel preparation. For the morphogen hub layer component, a commercially available solution of PEGDA 3.4K (Advanced Biomatrix, #GS700) was dissolved in UltraPure water to make a 30 wt% stock solution. Mebiol (Advanced Biomatrix, #5180), a component of the intermediate and organoid matrix layers, was prepared per the manufacturer’s instructions. Methacrylated hyaluronic acid was procured from Advanced Biomatrix as a kit with LAP (PhotoHA kit, #5274- 1KIT). PEGDA and M:P solutions were prepared at specified volumes to achieve a layer height of 1 mm each. The top M:H layer was prepared on ice as the last step for Models #1-#3 before photocrosslinking, and was added to give a total gel height of 5 mm. To polymerize, gels were first photo-crosslinked using 405 nm light exposure (10 mW/cm^2^) for 1 minute. Gels were left to rest on ice for 5 minutes, then transferred to a 37℃ incubator for 15 minutes. Following the thermal crosslinking for Mebiol-containing layers, neural induction medium (SAS CD, defined in Extended Methods, Supplementary Information) was added to hydrate gels. BMP4 (Gibco, PHC9534) and CHIR-99021 (Tocris Bioscience, 442310) were added to the M:H layer in Model #1 and the PEGDA layer in Models #2–#5. BMP4 at 400 ng/mL was supplemented with 24 µM CHIR-99021, while BMP4 at 4000 ng/mL was supplemented with 240 µM CHIR-99021. For hIPS2 iPSC, BMP4 at 200 ng/mL was supplemented with 24 µM CHIR-99021, while BMP4 at 2000 ng/mL was supplemented with 240 µM CHIR-99021.

### Digital light processing (DLP) printing of hydrogels (Models #4, #5, gradient, and two-morphogen system)

DLP-based 3-D printing was performed using the CELLINK BIONOVA X. All gels were printed at 25°C using a 24-well glass-bottom plate and a 24-well probe (CELLINK). Upon fabrication, gels were washed and stored in UltraPure water. Gels printed for rheology measurements were stored in UltraPure water overnight at 4°C before testing. The 24-well glass-bottom plates (Cellvis) used for hydrogel preparation were pre-treated to improve attachment at the glass-gel interface. For pre-treatment, the plate was first exposed to O2 plasma treatment for 10 minutes. A 10% 3-(trimethoxysilyl)propyl methacrylate (TMS, Thermo Scientific, 216555000) in ethanol solution was added to each well and left to coat for 1 hour. Following the TMS coating, treated wells were washed 5× with 100% ethanol. Prior to use, TMS-coated plates were exposed to UV for 30 minutes.

For printing of the microwell-containing hydrogel models, the morphogen hub layer (PEGDA) was printed first as a 6 mm diameter x 1 mm height cylinder and washed with PBS (Gibco). The intermediate layer (M:P) was printed over the morphogen hub with design parameters set as a 9 mm diameter x 4 mm height cylinder, with two 4 mm diameter ⅹ 1 mm height wells at the top (total gel height 5 mm). Microwells were coated to improve gel attachment by adding poly-L-lysine (Sigma-Aldrich, P4707) to the wells for 30 minutes. Wells were washed with PBS 3x and dried before adding the organoid gels.

### Modeling of morphogen diffusion using a fluorescent marker

For the one-morphogen hydrogels, FITC-dextran (Sigma-Aldrich, FD40S) was used to model BMP4 (dimeric) diffusion. Gels were prepared with respect to the proper protocol based on the condition, up to the media addition step. Gels were imaged using the Zeiss LSM 900 confocal microscope. Brightfield mode was used to identify the true bottom of the plate by focusing on a mark made on the bottom of the plate with a permanent marker and noting the focus plane as the base of the z-stack. Gels were imaged using a 2.5x magnification. A 4000 µm z-stack was imaged starting from the bottom of the plate, with a slice distance of 20 µm. The FITC channel was used to image the hydrogels. Gels were imaged on Days 0, 1, 3, and 6, with media being changed on Day 3 to reflect the experimental maintenance protocols. The well plate was stored in a humidified incubator at 37°C with 40% O₂ and 5% CO₂ to mimic experimental conditions. For the two-morphogen hydrogels, FITC-dextran and TRITC-dextran (Sigma-Aldrich, 73766) were used to model BMP4 and FGF8b diffusion, respectively. Gels were prepared with respect to the proper protocol based on condition (bulk vs. gradient stiffness). The imaging and maintenance protocols for the 1-morphogen hydrogels were used for the 2-morphogen hydrogels. The Rhodamine B channel was used to image TRITC. Gels were imaged on days 0, 1, 3, and 6, with a media change on day 3 to reflect the theoretical experimental protocol. Gels were stored in a humidified incubator at 37°C with 40% O₂ and 5% CO₂ to mimic experimental conditions. Details for the fluorescence image analysis for all hydrogels can be found in the Supplementary Information.

### Rheological measurements

Rheological measurements were performed using a TA DHR-3 rheometer. An 8-mm cross-hatched plate (TA Instruments) was used for the upper geometry and a standard Peltier plate was used for the lower plate. PEGDA and M:P samples were printed and stored in PBS overnight at 37°C before measurements were conducted. M:H gels were prepared following the experimental protocol. All experiments were performed at a temperature of 37℃. Amplitude sweeps were performed at 1 Hz from 0.01-500% strain with 5 points/decade. Frequency sweeps were performed at 0.1% strain from 0.1– ∼30 Hz with 5 points/decade. For frequency sweeps, samples were enclosed in a humidity chamber during testing to minimize solvent evaporation.

### Stem cell culture

All experiments involving human pluripotent stem cells were approved by the Human Stem Cell Research Oversight (hSCRO) committee at the University of California, Irvine. The H9 human embryonic stem cell (hESC) line was obtained from the WiCell Institute, and the XF and hIPS2 human induced pluripotent stem cell (hiPSC) lines were obtained from the UCLA Broad Stem Cell Research Center Core under a material transfer agreement. The H9 hESC and hIPS2 hiPSC lines were used for all primary experiments, whereas the XF hiPSC line was used for the compatibility assays. Stem cells were cultured with mitotically inactivated mouse embryonic fibroblast (MEF) feeder layers, including mitomycin C–treated MEFs (PMEF-CF) (MilliporeSigma), on 0.1% gelatin-coated tissue culture plates. Cells were maintained in hESC medium composed of DMEM/F12 (Hyclone, SH30023.02), 20% KnockOut Serum Replacement (KSR; Life Technologies, 10828-028), non-essential amino acids (NEAA, 100×; Life Technologies, 11140-050), GlutaMAX (100×; Life Technologies, 35050-061), 100 µg/mL Primocin (InvivoGen, ant-pm-2 or ant-pm-1), 0.1 mM β-mercaptoethanol (Gibco, 21985-023), and 10 ng/mL fibroblast growth factor 2 (FGF2; Invitrogen, PHG0021).

The culture medium was refreshed daily. Cells were maintained at 37°C in a humidified incubator with 5% CO₂. Stem cells were passaged every six days using the StemPro EZ Passage Tool (Invitrogen) and replated at a 1:4 to 1:8 split ratio depending on confluency. Details for the maintenance of organoid-seeded hydrogels, as well as quantitative PCR and immunohistochemistry analyses for organoids, can be found in the Supplementary Information.

### Organoid seeding for bulk hydrogel experiments

Bulk and DLP-printed gels were prepared as described above. All gels were prepared in a 24-well glass-bottom plate (Cellvis). Organoids in culture medium were added directly into the liquid-state gel precursor for the organoid layer of the bulk hydrogels. The organoid layer was crosslinked using 405 nm light exposure (10 mW/cm^2^, 1 min). Following this, the plate was left to rest on ice for 5 min, followed by incubation at 37℃ for 15 min. After incubation, 200 µL of SAS CD media were added to each gel. For the DLP-printed gels, organoids were first mixed with the organoid gel formulation to generate a uniform suspension, which was then seeded into the microwells. An additional layer of gel precursor was subsequently added to improve adhesion between the organoid-containing layer and the microwell surface. The organoid layer was polymerized through a multi-step series, starting with 405 nm light exposure (10 mW/cm^2^) for 1 min. Gels were left to rest on ice for 5 min and transferred to 37°C for 15 min. After gelation, 1000 µL of SAS-CD media were added to fully submerge gels. Embedded organoids were maintained in a humidified incubator at 37°C with 40% O₂ and 5% CO2.

### Confocal microscopy for organoid characterization

Images were collected (2048 × 2048 px) with a 10× objective using the Zeiss LSM 900 Airyscan 2 (Zen Blue 3.3 software) mode. The same acquisition settings were used within the experimental sets for comparisons between groups. Post-acquisition adjustments, such as brightness and contrast, were performed using ImageJ/Fiji and Adobe Photoshop, and the same adjustment settings were uniformly applied to all groups from the same experimental set. For organoid characterization, cell number quantification was performed manually by using the Cell Counter function in ImageJ/Fiji. The percentage of cells was then calculated by normalizing to the total number of live cells.

## Supporting information

Supplementary Information, revised as of 10/2026

## Acknowledgements

This project has been supported by the NSF-CBET RECODE 2225624, New Investigator Awards from the UCI School of Medicine (M.W.); FRAXA Postdoctoral Fellowship (Y.C.T.); postdoctoral training grant from the California Institute for Regenerative Medicine (CIRM), EDUC4-12822 (H.O.), and UCI start-up funds (H.A.M.A.). The authors acknowledge the use of facilities and instrumentation at the UC Irvine Materials Research Institute (IMRI), which is supported in part by the National Science Foundation through the UCI Materials Research Science and Engineering Center (DMR-2011967). We also appreciate the support of the UC Irvine CIRM Shared Resources Laboratories to Enhance In Vitro Stem Cell Modeling and Training Grant INFR6.2-15368.

## Data Availability Statement

The data that support the findings of this study are available from the corresponding author upon reasonable request.

