## Supplementary Information, revised as of 10/2026 for "Spatial engineering of posterior signaling centers in cerebral organoids via controlled morphogen exposure within hydrogels"

### Extended Methods

*Fluorescence image analysis.* Fluorescence Z-stack images (.tiff) were extracted from the .czi files, then analyzed using a custom Python-based workflow. For one-morphogen hub models (Models #1, #2, #3, #4, and gradient hydrogels), the FITC channel was extracted as a 3-D image volume. All FITC stacks were scanned to define a shared FITC range, with low boundary intensity ($I_{low}$) set to the minimum 0.5th percentile and high boundary intensity ($I_{high}$) set to the maximum 99.5th percentile across all analyzed volumes. To avoid per-image contrast rescaling, raw fluorescence intensities were normalized using global percentile-based intensity ranges. Each image volume was then rescaled based on percentile-based normalization (0 to 1) as:

$$I_{norm} = \frac{I_{raw} - I_{low}}{I_{high} - I_{low}}$$

For two-morphogen hub models, the same procedure was applied independently to FITC-dextran and TRITC-dextran, yielding channel-specific global normalization ranges. Normalized values are reported as arbitrary fluorescence units and were used for both cross-sectional visualization and Z-layer mean intensity profiles.

Axial signal distribution was assessed by calculating mean fluorescence intensity across the XY plane for each Z layer. Both transverse and coronal cross-sectional heatmaps were generated from normalized image volumes at selected Z-depths using fixed display intensity ranges across comparable models. For one-morphogen hub models, coronal XZ sections were extracted at the middle Y plane to visualize FITC-dextran distribution across depth and time. For two-morphogen hub models, FITC-dextran and TRITC-dextran were displayed in matched cross-sectional layouts, and diagonal coronal sections were extracted along the axis connecting the FITC- and TRITC-loaded regions to visualize relative channel distribution. Gaussian smoothing and color-scale adjustment were applied only for visualization and did not alter the underlying Z-layer intensity profiles.

*Enzyme-linked immunosorbent assay (ELISA).* Hydrogel Models #1–3 were generated as described in the Materials and Methods, with four experimental replicates per model. Following thermal crosslinking of the Mebiol-containing layers, 200 µL of Dulbecco's phosphate-buffered saline (DPBS) was added to each well to hydrate the gels. After incubation at 37°C for 24 h, DPBS was collected from each well for BMP-4 quantification using the Human BMP-4 ELISA Kit (Sigma-Aldrich, RAB0030) according to the manufacturer's protocol. Briefly, samples were diluted fivefold, and 100 µL of each standard or diluted sample was added to the ELISA plate in technical duplicate. The plate was incubated for 2.5 h at room temperature with gentle shaking, followed by four washes with wash buffer. Detection antibody (100 µL per well) was then added, and the plate was incubated for 1 h at room temperature with gentle shaking. After four washes, streptavidin solution (100 µL per well) was added, followed by incubation for 45 min at room temperature with gentle shaking. The plate was washed four times, and colorimetric reagent (100 µL per well) was added and incubated for 30 min at room temperature in the dark with gentle shaking. Stop solution (50 µL per well) was then added, and absorbance at 450 nm was immediately measured using a BioTek Cytation 5 plate reader (Agilent).

*Development of solutions for morphogen diffusion simulations.* To quantitatively assess the ability of the layered hydrogel configurations to delay morphogen transport, an unsteady mass transfer model was developed to monitor the evolution of morphogen concentration profiles within the hydrogel constructs. Assuming one-dimensional diffusion in the z-direction and negligible morphogen consumption or reaction within the hydrogels, transport was governed by Fick's second law:

$$\frac{\partial C_i}{\partial t} = D_i \frac{\partial^2 C_i}{\partial z^2}$$

where $C_i$ is the morphogen concentration and $D_i$ is the diffusivity within medium $i$, corresponding to the different hydrogel layers used to modulate transport. The initial condition consisted of a uniform morphogen concentration within the morphogen hub layer and zero concentration elsewhere in the construct. No-flux boundary conditions were imposed at the external boundaries of the domain.

To generalize the transport behavior across the different hydrogel scenarios, the governing equations were non-dimensionalized using:

$$\theta = \frac{C_i}{C_0},\ \eta = \frac{z}{L},\ \tau = \frac{tD_{ref}}{L^2},$$

where $C_0$ is the initial morphogen concentration in the morphogen hub, $L$ is the total hydrogel height, and $D_{ref}$ corresponds to the diffusivity of the softest hydrogel layer, which exhibits the fastest characteristic diffusion timescale. This reference condition corresponds to Scenario #1 in Figure 3, where the construct consists entirely of the softest material. Consequently, comparisons between the different modeled scenarios only require knowledge of the relative diffusivities between the hydrogel layers. Relative diffusivities were approximated as inversely proportional to the experimentally measured stiffness ratios obtained from rheological characterization.

The resulting non-dimensional equations were solved numerically using second-order finite difference approximations for the spatial derivatives and explicit Euler integration for the temporal evolution. Spatial and temporal discretizations were selected as $\Delta\eta = 2 \times 10^{-3}$ and $\Delta\tau = 2 \times 10^{-7}$ to ensure numerical stability across all simulated conditions.

For multi-layered systems containing interfaces between materials of different diffusivities (Scenarios #2–#4), continuity of both concentration and diffusive flux was imposed at each interface. At the discretized interface node, the effective diffusivity was modified to preserve interfacial flux continuity between adjacent media. The simulations were used to generate concentration profiles throughout the hydrogel thickness at multiple diffusion timescales, as well as temporal concentration evolution at the organoid location ($\eta = 0.8$), corresponding to the approximate position where the diffusing morphogen front interfaces with the organoid layer.

*Cortical organoid generation and maintenance.* Cortical organoids were generated as previously described.[1-3] Briefly, once hESC or hiPSC colonies reached 70–80% confluency, they were detached using dispase (STEMCELL Technologies, 7913) and dissociated with 0.05% trypsin–EDTA (Gibco, 25300-054) supplemented with DNase I (Worthington, LK003172). Cell clumps were removed by passing the suspension through a 100 μm cell strainer. A total of 9,000 cells per well were seeded into low-attachment V-bottom 96-well plates (Sumitomo Bakelite, MS9096V) and aggregated by centrifugation at 290 × g for 2 min. From day (D) 0 to D15, organoids were cultured in neural induction medium (hereafter referred to as SASAI Cortical Differentiation (SAS CD) medium) consisting of Glasgow's Minimum Essential Medium (GMEM; Gibco, 11-710-035), 20% KnockOut Serum Replacement (KSR; Gibco, 10828-028), non-essential amino acids (NEAA), 100 μg/mL Primocin, 0.1 mM β-mercaptoethanol, and 1 mM sodium pyruvate (Gibco, 11-360-070). To promote cortical fate specification, the WNT signaling inhibitor IWR-1-endo (IWR1e; Calbiochem, 681669) and the TGF-β signaling inhibitor SB431542 (SB; Stemgent, 04-

0010-103) were added to SAS CD medium. The concentrations of these inhibitors were empirically optimized for each cell line. H9 hESCs were treated with 3 μM IWR1e and 3 μM SB, whereas XF hiPSCs were treated with 1 μM IWR1e and 1 μM SB. In addition, 20 μM of the ROCK inhibitor Y-27632 (BioPioneer, SM-008) was supplemented during the first six days of differentiation (D0–6) to enhance cell survival. Half of the culture medium was refreshed every three days until D15. Organoids were maintained at 37°C in a humidified incubator with 5% $CO_2$. For the posterior signaling center induction experiments, Brilliant Blue G (Tokyo Chemical Industry) was introduced to the unexposed side of each organoid upon retrieval from the hydrogels.

*Maintenance of organoid-seeded hydrogels.* Embedded organoids were maintained at 37°C for 6 days (up to Day 21 for organoids). The culture media were refreshed daily. On day 3 (D18 for organoids), media volume was increased to 400 uL to support continued organoid growth. On day 6 (Day 21 for organoids), organoids were extracted from the hydrogel layer, transferred to Petri dishes containing N2 medium, and maintained until Day 35, consisting of DMEM/F12 (Hyclone, SH30023.02), N2 supplement (100×; Life Technologies, 17502-048), chemically defined lipid concentrate (CDLC, 100×; Life Technologies, 11905-031), GlutaMAX, and 100 μg/mL Primocin, and 0.4% methylcellulose (Sigma, M7140-100G).

*Quantitative PCR.* Organoids were collected at Day 35, and 4–6 organoids were pooled to generate one biological sample. Samples were lysed, and total RNA was extracted using the RNeasy Mini Kit (Qiagen, 79216) according to the manufacturer's instructions. For cDNA synthesis, 1,000–2,500 ng of total RNA per sample was reverse-transcribed using the

SuperScript™ IV First-Strand Synthesis System (Invitrogen, 18091050). Real-time quantitative PCR was performed using PowerTrack SYBR Green Master Mix (Applied Biosystems, A46109), gene-specific primer pairs (Integrated DNA Technologies), and 30 ng of cDNA per reaction. For each gene of interest, at least three biological replicates and technical replicates were included. Reactions were loaded into 384-well plates (Invitrogen, 4309849) and run on a QuantStudio 6 or 7 Real-Time PCR System (Applied Biosystems). Relative gene expression levels were calculated using the ΔCt method and normalized to GAPDH expression. Primers used in this study are listed below. Statistical analysis was performed using one-way ANOVA using GraphPad Prism. Data are presented as mean ± standard deviation (SD).

*Primer list:*

| **Gene** | **Forward** | **Reverse** |
|---|---|---|
| COUPTF1 | ATCGTGCTGTTCACGTCAGAC | TGGCTCCTCACGTACTCCTC |
| MSX1 | AGTTCTCCAGCTCGCTCAGC | GGAACCATATCTTCACCTGCGT |
| OTX2 | AGAGGACGACGTTCACTCG | TCGGGCAAGTTGATTTTCAGT |
| SP8 | CTTCACTTCTAGGGGAAGAACC | AGCTTGAGAGACTGGAACCC |
| TTR | ATCCAAGTGTCCTCTGATGGT | GCCAAGTGCCTTCCAGTAAGA |

*Immunohistochemistry.* Organoids were collected at Day 35 and fixed in 4% paraformaldehyde in phosphate-buffered saline (PBS) for 20 min on ice. Samples were incubated in 30% sucrose in PBS on ice for cryoprotection, then embedded in Tissue-Tek Optimal Cutting Temperature (OCT) compound (Sakura, 25608-930) using cryomolds (Sakura, 25608-924).

Embedded organoids were cryosectioned at a thickness of 12 µm using a cryostat (Leica, CM3050S) and mounted onto Superfrost™ Plus microscope slides (Fisherbrand, 12-550-15).

Immunostaining was performed as previously described.[4] Briefly, organoid sections were incubated in blocking buffer consisting of 0.1% heat-inactivated horse serum (donor equine serum; Hyclone, SH30074.02), 0.1% Triton X-100, 0.01% sodium azide, and PBS for 1 h at room temperature. Primary antibodies (below) were diluted in blocking buffer and added to the sections overnight at 4°C. Sections were then washed three times with PBS containing 0.1% Triton X-100 (PBST), followed by incubation with appropriate secondary antibodies and Hoechst 33258 for 1 h at room temperature. After additional washes with PBST, sections were coverslipped using ProLong™ Diamond Antifade Mountant (Life Technologies, P36970).

Cell counts were analyzed using one-way ANOVA using GraphPad Prism. Data are presented as mean ± SD. For immunofluorescence intensity analyses, fluorescence intensity profiles were measured across the width of representative regions of the organoid sections, with start and end points selected to cover the full thickness of the region of interest. The distance axis was normalized to the total distance measured and scaled from 0 to 1, and the profiles are presented as mean ± SD.

*Antibody list:*

| **Protein** | **Species** | **Company** | **Catalog#** | **Dilution** |
|---|---|---|---|---|
| cl-CASP3 | Rabbit | Cell Signaling | 9961S | 1:500 |
| OTX2 | Rabbit | Abcam | ab21990 | 1:1000 |
| PAX6 | Mouse | DSHB | N/A | 1:100 |
| TTR | Rabbit | Dako | A000202-2 | 1:1000 |

### Supplementary Data

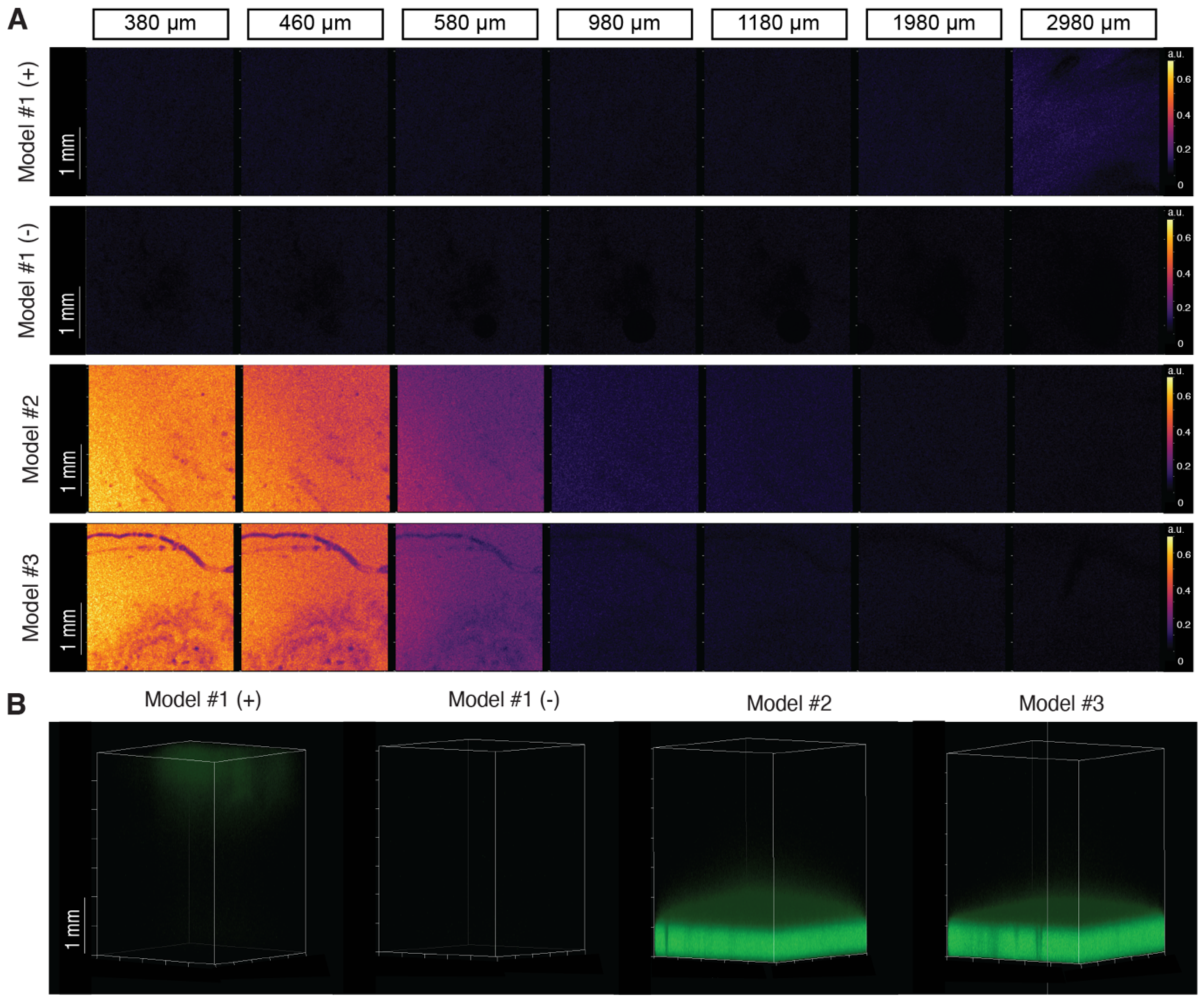


**Figure S1. Visualization of varied morphogen diffusion profiles for Models #1–#3 at Day 0.** (A) Representative diffusion heatmaps across different transverse planes (Z= 380, 460, 580, 980, 1180, 1980, and 2980 µm). (B) 3-D reconstruction of FITC-dextran fluorescence levels. Data represent Models #1 (0 and 400 ng/mL, negative and positive controls, respectively), #2, and #3. Scale bars: 1 mm.

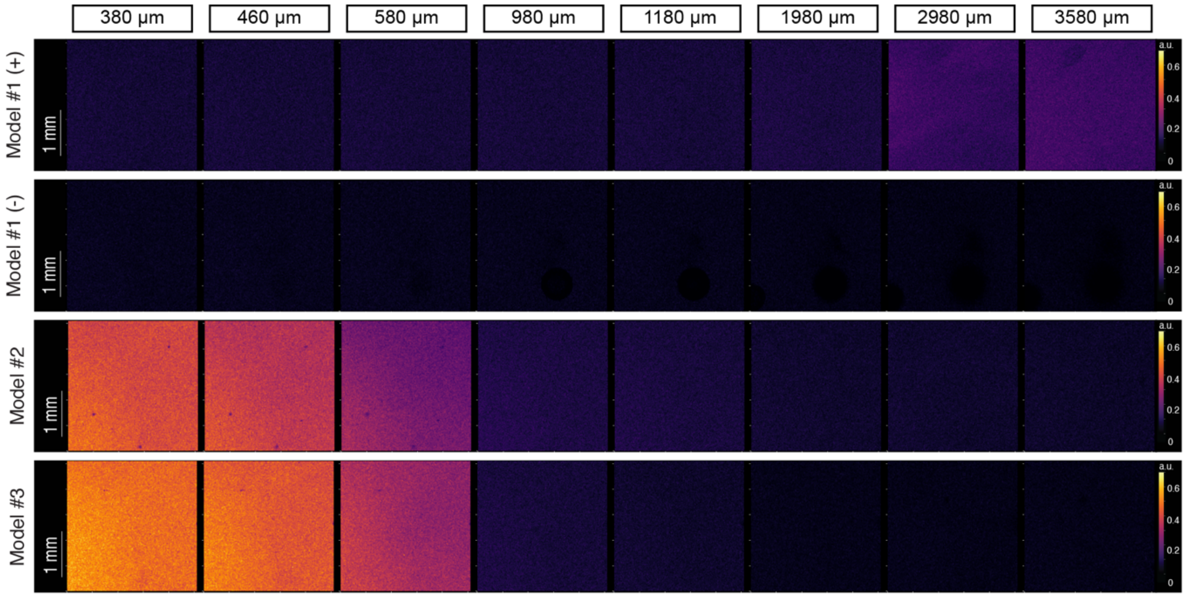


**Figure S2. Visualization of varied morphogen diffusion profiles for Models #1–#3 at Day 1.** Representative diffusion heatmaps across different transverse planes (Z= 380, 460, 580, 980, 1180, 1980, 2980, and 3580 µm). Data represent Models #1 (0 and 400 ng/mL, negative and positive controls, respectively), #2, and #3. Scale bar: 1 mm.

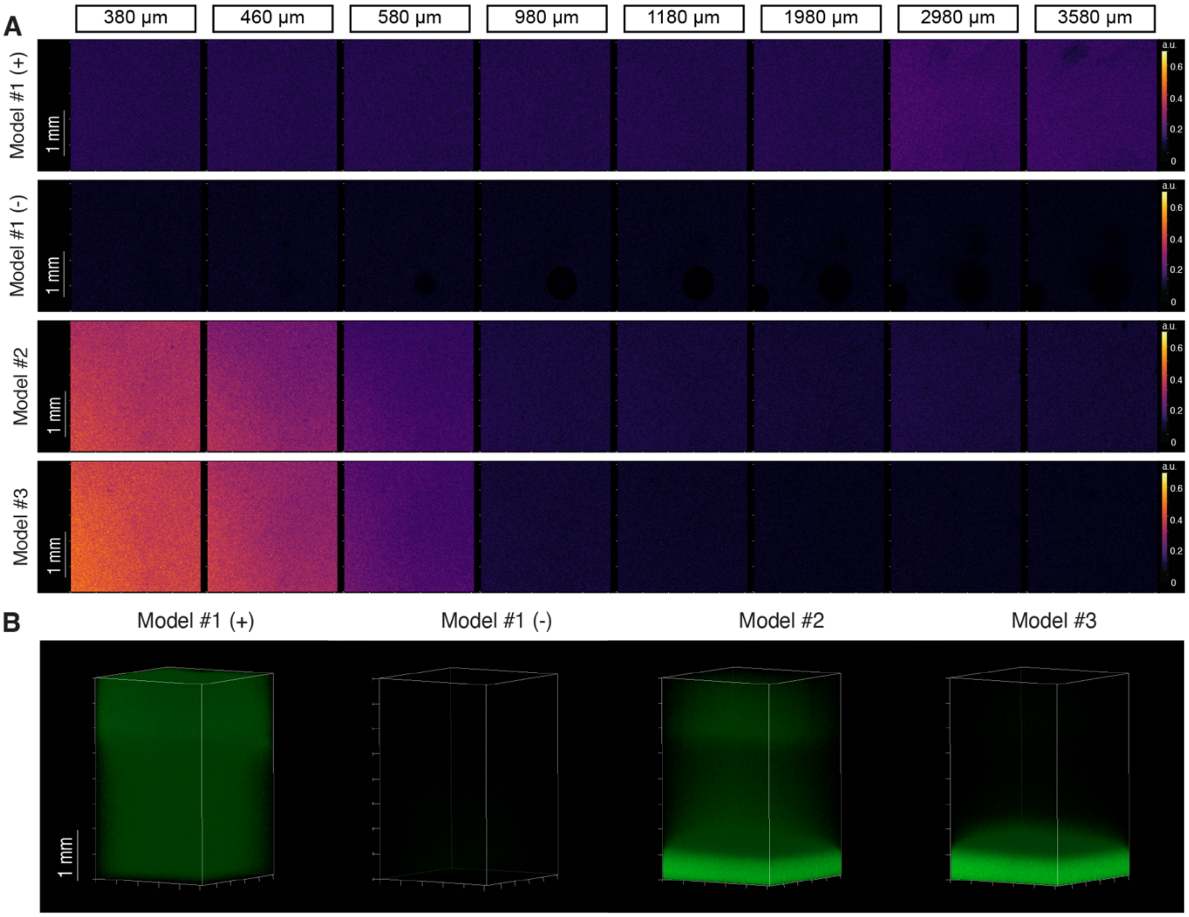


**Figure S3. Visualization of varied morphogen diffusion profiles for Models #1–#3 at Day 3.** (A) Representative diffusion heatmaps across different transverse planes (Z= 380, 460, 580, 980, 1180, 1980, 2980, and 3580 µm). (B) 3-D reconstruction of FITC-dextran fluorescence levels. Data represent Models #1 (0 and 400 ng/mL, negative and positive controls, respectively), #2, and #3. Scale bars: 1 mm.

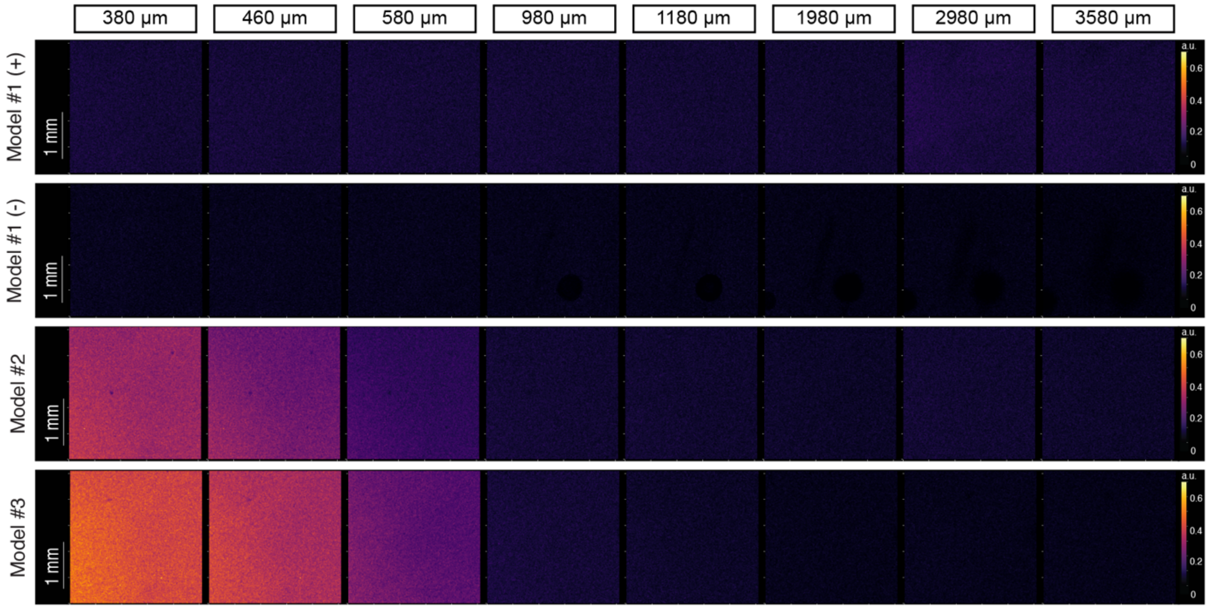


**Figure S4. Visualization of varied morphogen diffusion profiles for Models #1–#3 at Day 6.** Representative diffusion heatmaps across different transverse planes (Z= 380, 460, 580, 980, 1180, 1980, 2980, and 3580 µm). Data represent Models #1 (0 and 400 ng/mL, negative and positive controls, respectively), #2, and #3. Scale bar: 1 mm.

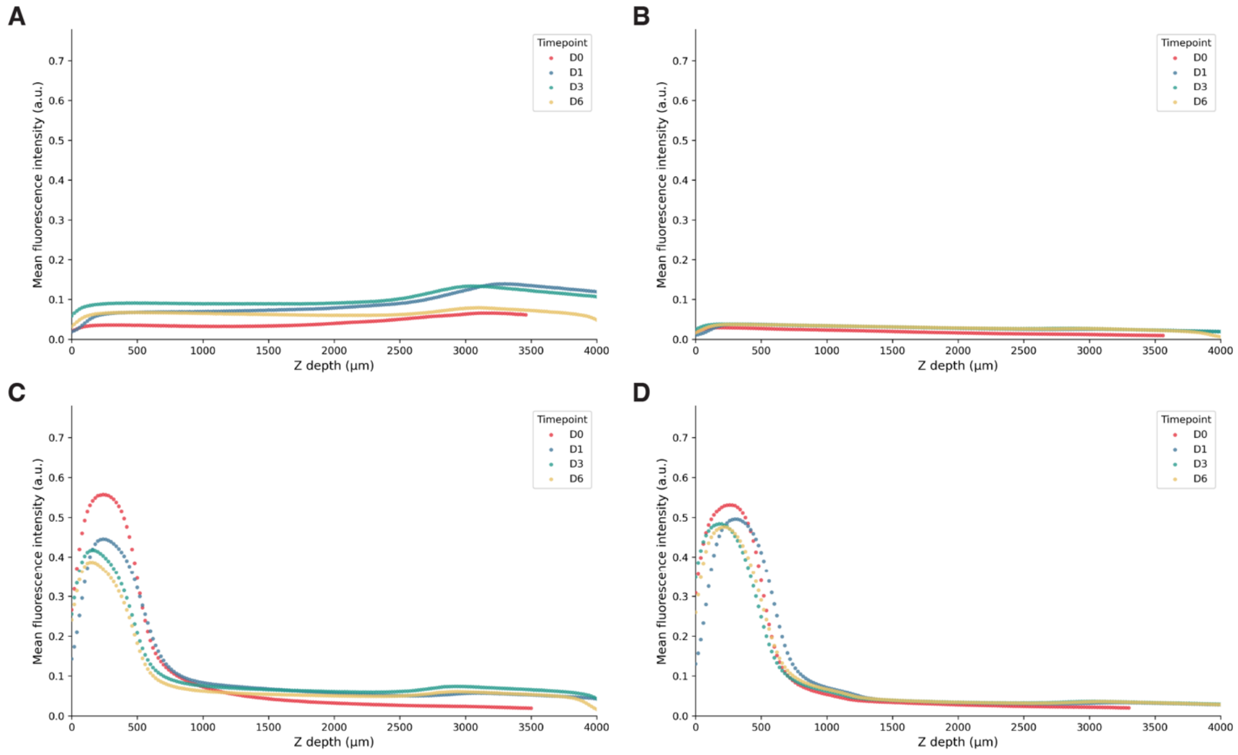


**Figure S5. Mean fluorescence intensity profiles across the Z-axis.** Day 0 to Day 6 mean fluorescence intensity Z-profiles are shown for: (A) Model #1 (+, 400 ng/mL); (B) Model #1 (-, 0 ng/mL); (C) Model #2; and (D) Model #3. These profiles are extracted from the same representative hydrogels analyzed for Figures 2 and S1–S4.

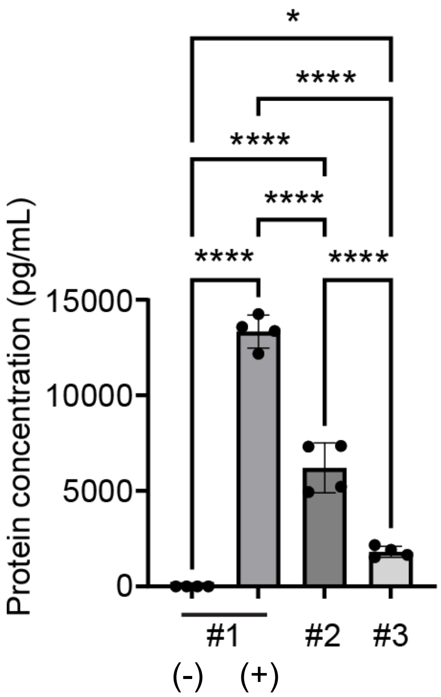


**Figure S6. BMP4 ELISA for Models #1–3**. BMP4 protein concentrations were measured in DPBS from Models #1–3 (for #1: 0 and 200 ng/mL BMP4, (-) negative and (+) positive controls, respectively; for #2 and #3, 2000 ng/mL), 24 hours after incubation. Four experimental replicates per model. Statistics: One-way ANOVA, Tukey's multiple comparisons test. Data are presented as mean ± standard deviation. *$p<0.05$; ****$p<0.0001$.

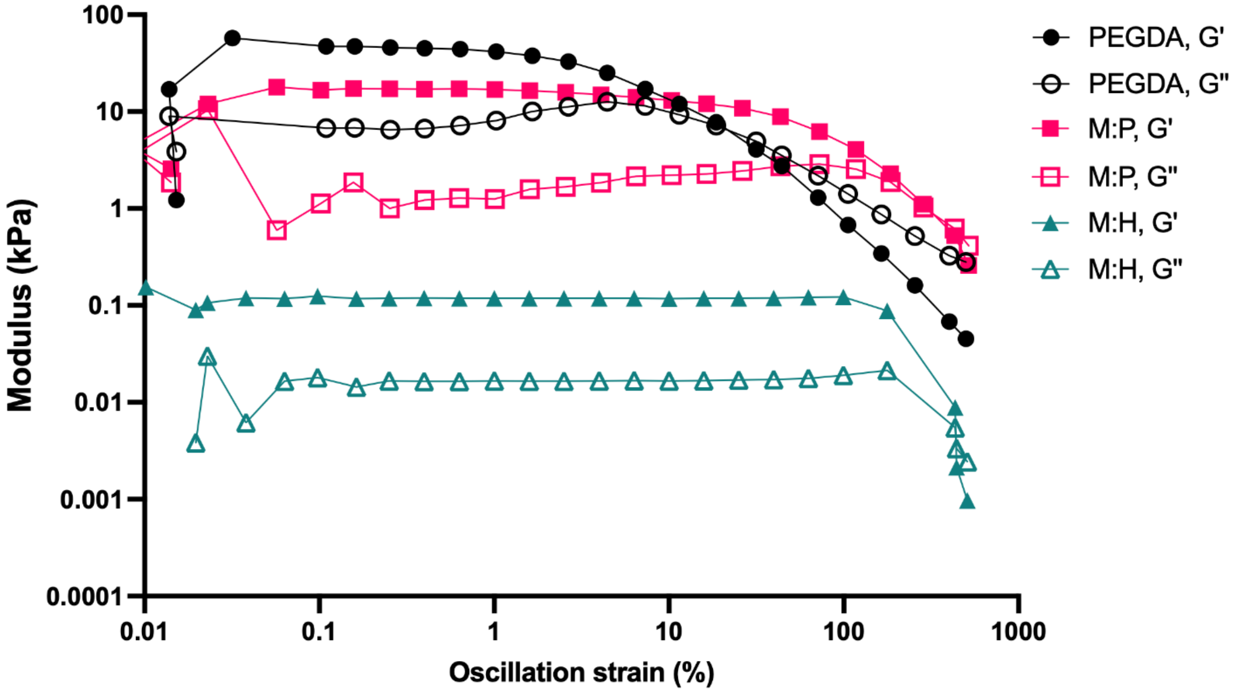


**Figure S7. Amplitude sweeps for the material library.** Representative amplitude (oscillation strain) sweeps for all polymeric formulations used in hydrogel constructs studied here. Measurements were all taken at 37°C. M:P and PEGDA shown were printed using 60% light intensity using the DLP printer setup used for Model #4 and #5. M:H was prepared under standard procedure of 405 nm photoirradiation (10 mW/cm$^2$) for 1 min, followed by 5 min rest, and 15 min incubation at 37°C.

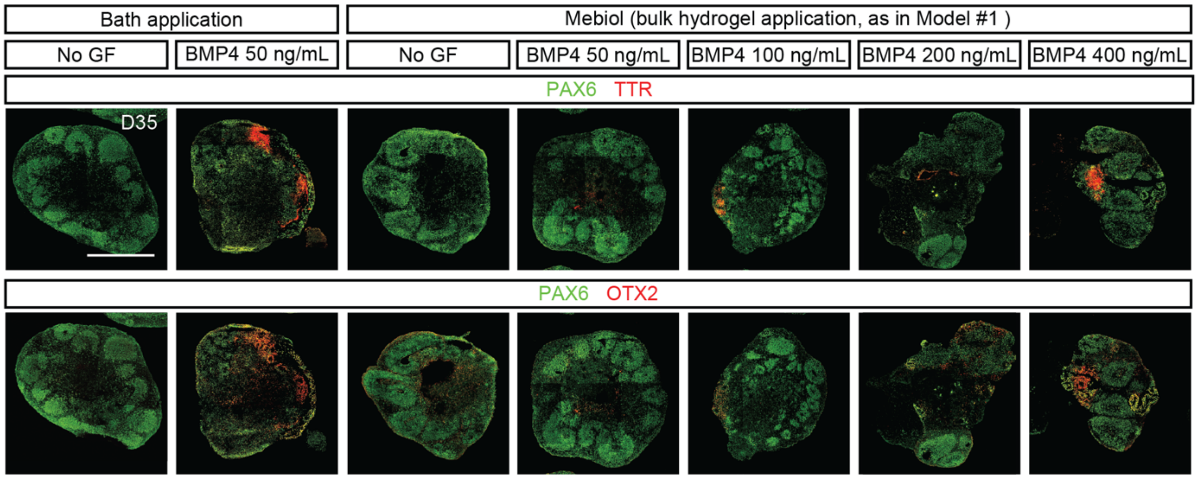


**Figure S8. Morphogen titration in bulk hydrogel loading method.** Data for morphogen delivery via bath application is added as a reference. Scale bar: 500 μm.

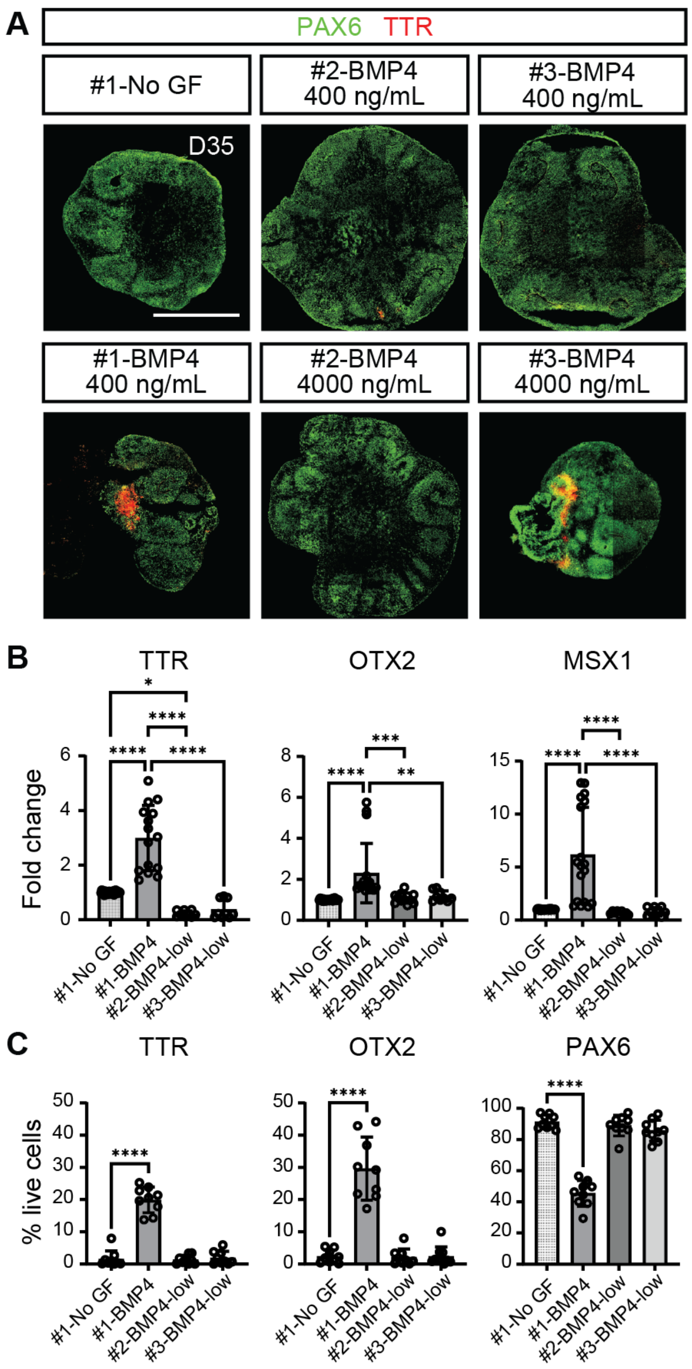


**Figure S9. Additional representative images for the immunochemical analyses of organoids under Models #1–#3 hydrogel conditions.** (A) Day 35 organoids were subjected to immunochemical analyses for cortical (PAX6) and posterior signaling center (TTR) markers. Scale bar: 500 µm. (B) Quantification of posterior marker induction by RT-qPCR (N= 12–15 samples per condition, derived from 4–5 pooled organoids per sample, 3–4 independent batches). BMP-low: BMP4 400ng/mL for H9. (C) % positive cells out of total live cells quantified from the immunofluorescence images in A (N= 9 organoids per condition, 3–5 independent batches).

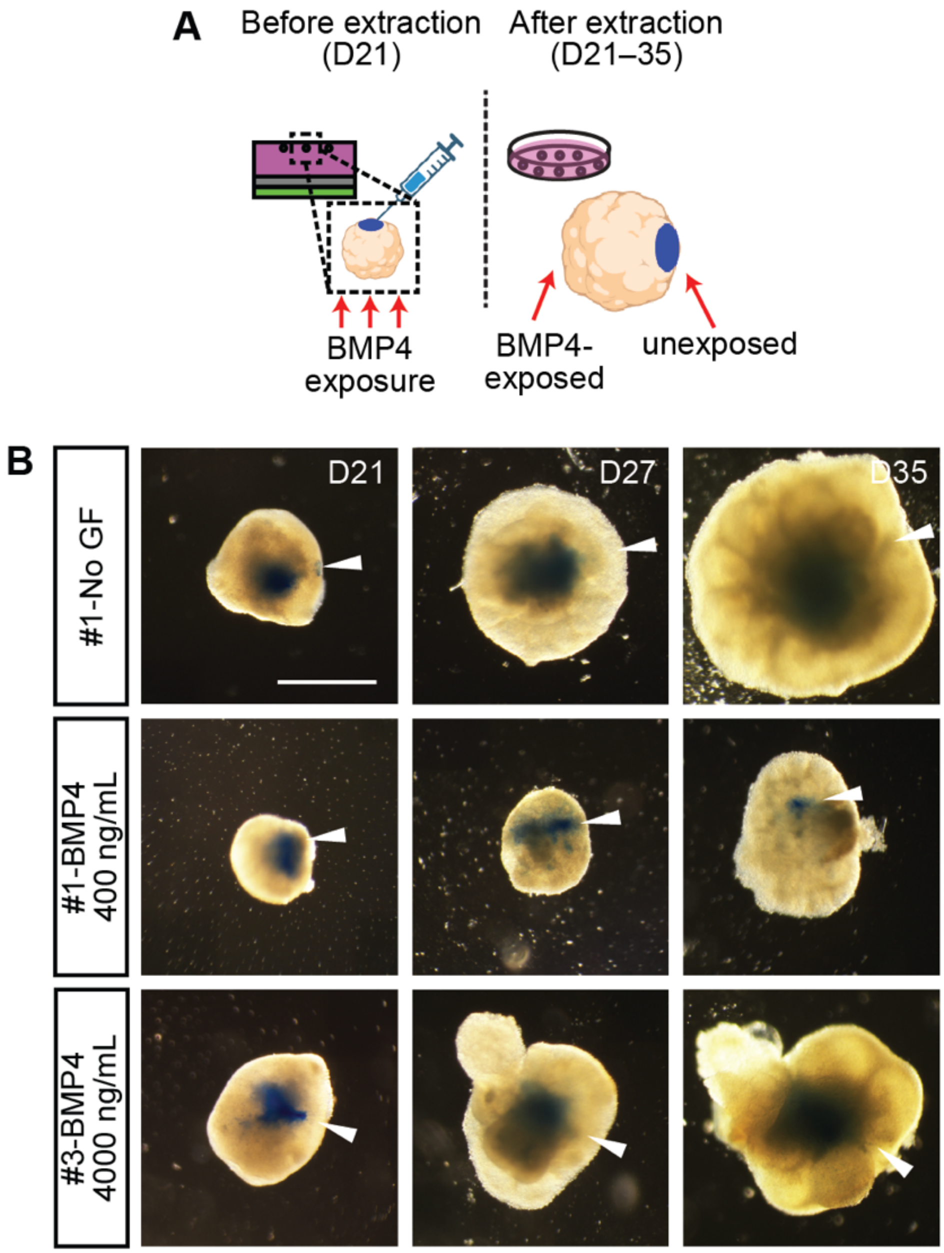


**Figure S10. Dye marking experiment for BMP4-exposed organoids.** (A) Schematic illustration of organoid marking with Brilliant Blue G dye. (B) Representative images of the blue dye-marked organoids. Scale bar: 500µm.

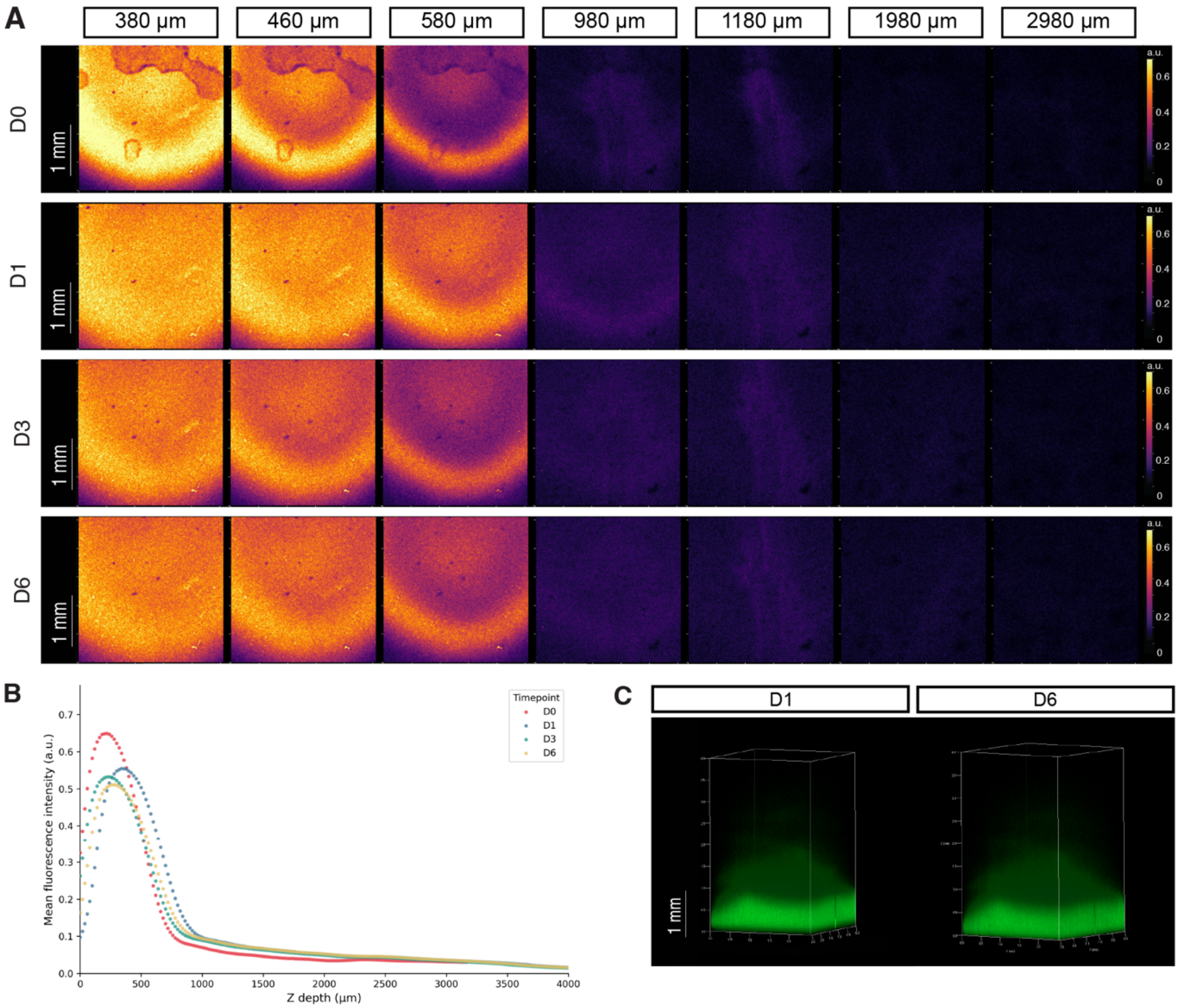


**Figure S11. Visualization of morphogen diffusion profiles for Model #4 hydrogels.** (A) Representative diffusion heatmaps across different transverse planes (Z= 380, 460, 580, 980, 1180, 1980, and 2980 µm). (B) Day 0 to Day 6 mean fluorescence intensity profiles across the Z-axis. 3-D reconstruction of FITC-dextran fluorescence levels at Day 1 and Day 6. Scale bars: 1 mm.

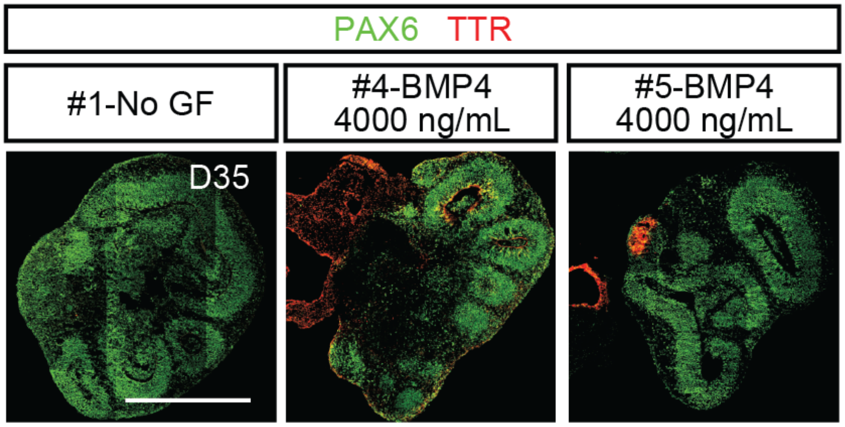


**Figure S12. Additional representative images for the immunochemical analyses of organoids under DLP-printed hydrogel conditions (Models #4 and #5).** Representative data from Model#1 was added as a reference. Day 35 organoids were subjected to immunochemical analyses for cortical (PAX6) and posterior signaling center (TTR) markers. Scale bar: 500 µm.

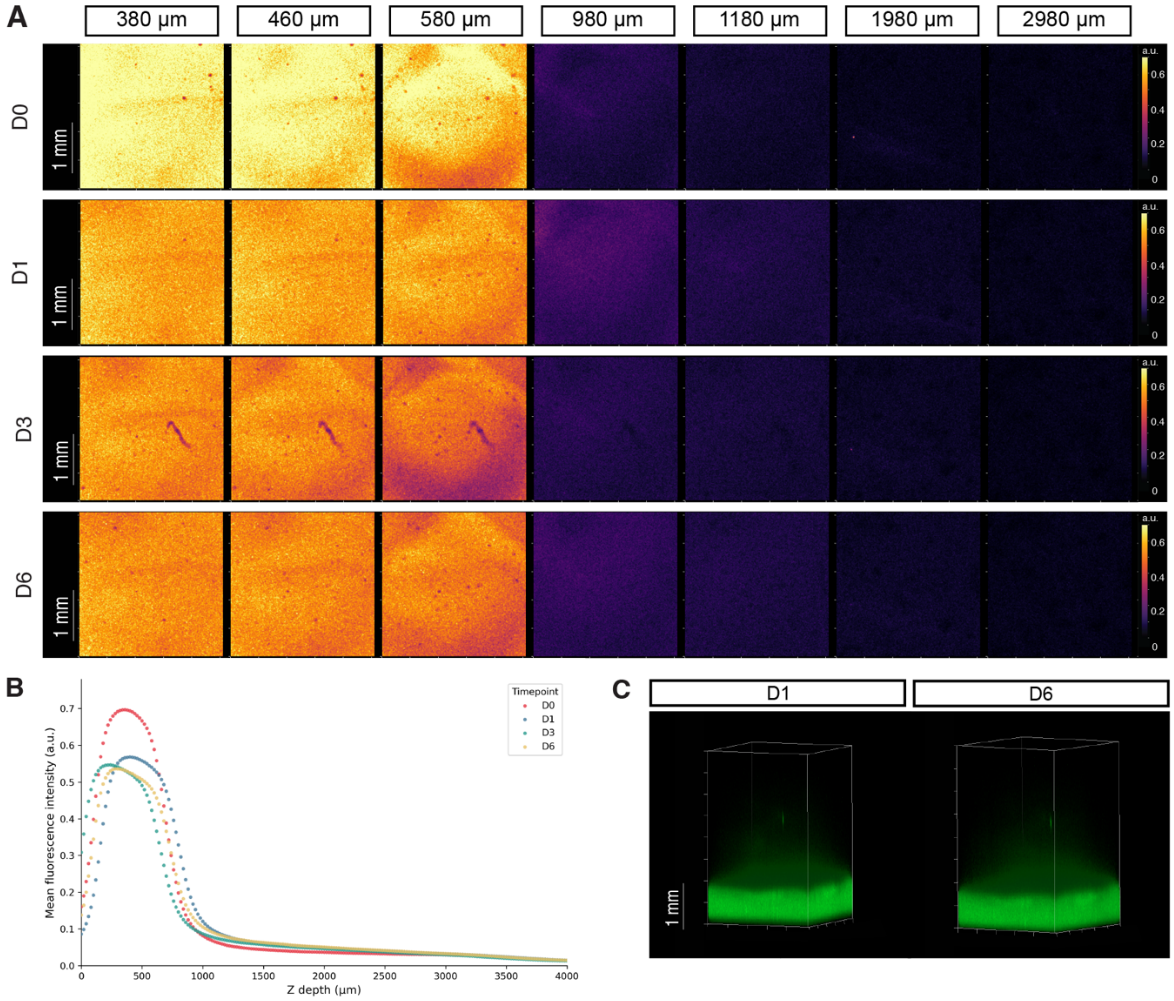


**Figure S13. Visualization of morphogen diffusion profiles for Model #5 hydrogels with stiffness gradients.** (A) Representative diffusion heatmaps across different transverse planes (Z= 380, 460, 580, 980, 1180, 1980, and 2980 µm). (B) Day 0 to Day 6 mean fluorescence intensity profiles across the Z-axis. 3-D reconstruction of FITC-dextran fluorescence levels at Day 1 and Day 6. Scale bars: 1 mm.

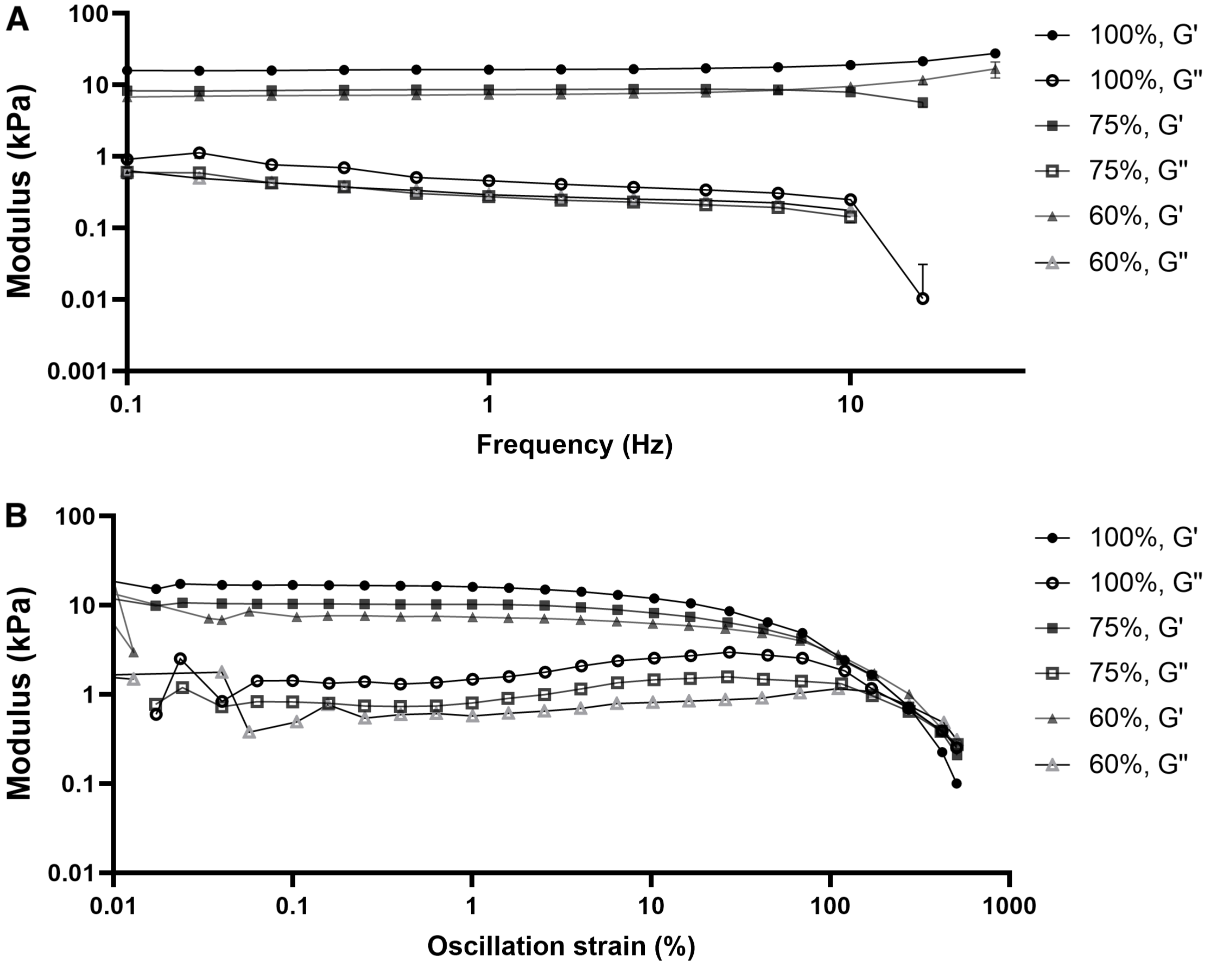


**Figure S14. Rheological characterization of hydrogels subjected to greyscale photomasking.** (A) Frequency sweep measurements at 0.1% strain at 37°C for the hydrogels used in modeling experiments in Figure 6 (N= 3; error bars represent standard error of the mean, s.e.m.). (B) Representative amplitude (oscillation strain) sweeps for the hydrogels processed in (A).

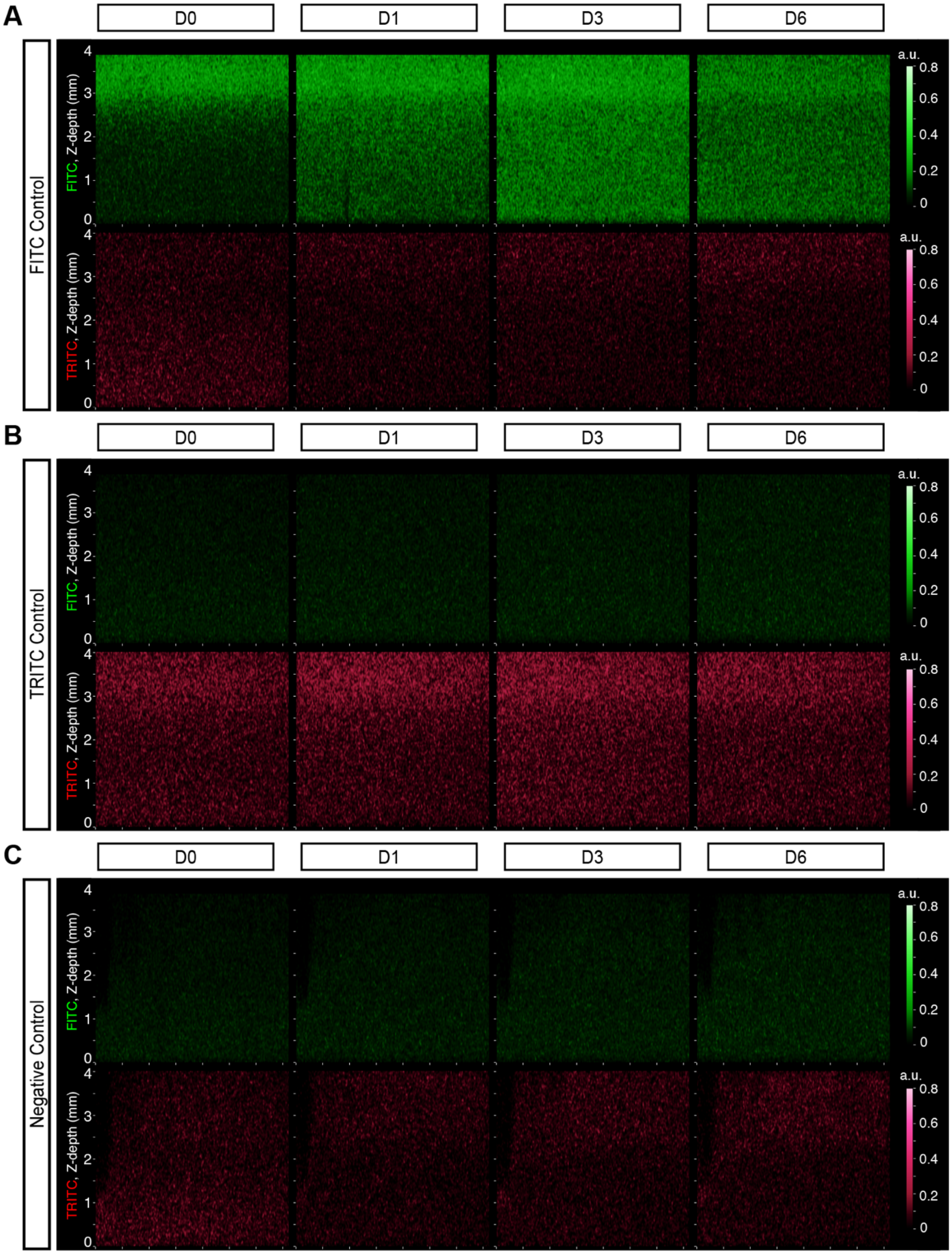


**Figure S15. Representative coronal plane cross-sections at the center of two-morphogen hydrogel controls.** Bulk loading positive controls for (a) FITC-dextran and (b) TRITC-dextran. No dextrans were added in the (c) negative control. Both FITC and TRITC channels were shown for data across Day 0 to Day 6.

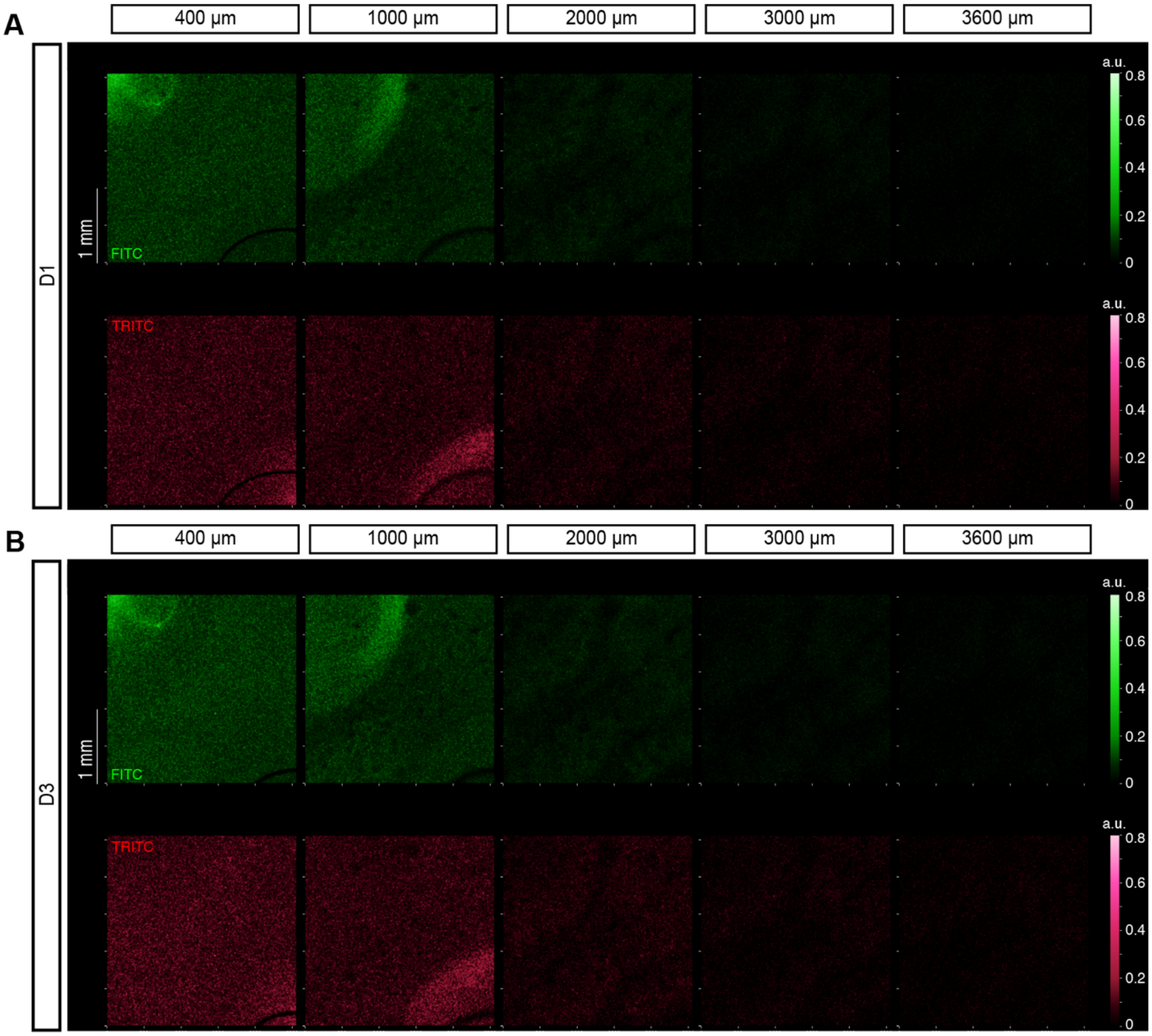


**Figure S16. Representative fluorescence images across different transverse planes for two-morphogen hydrogel constructs.** Data for (A) Day 1 and (B) Day 3 at Z= 400, 1000, 2000, 3000, and 3600 μm are shown. These images were taken from the same hydrogels presented in Figure 6.

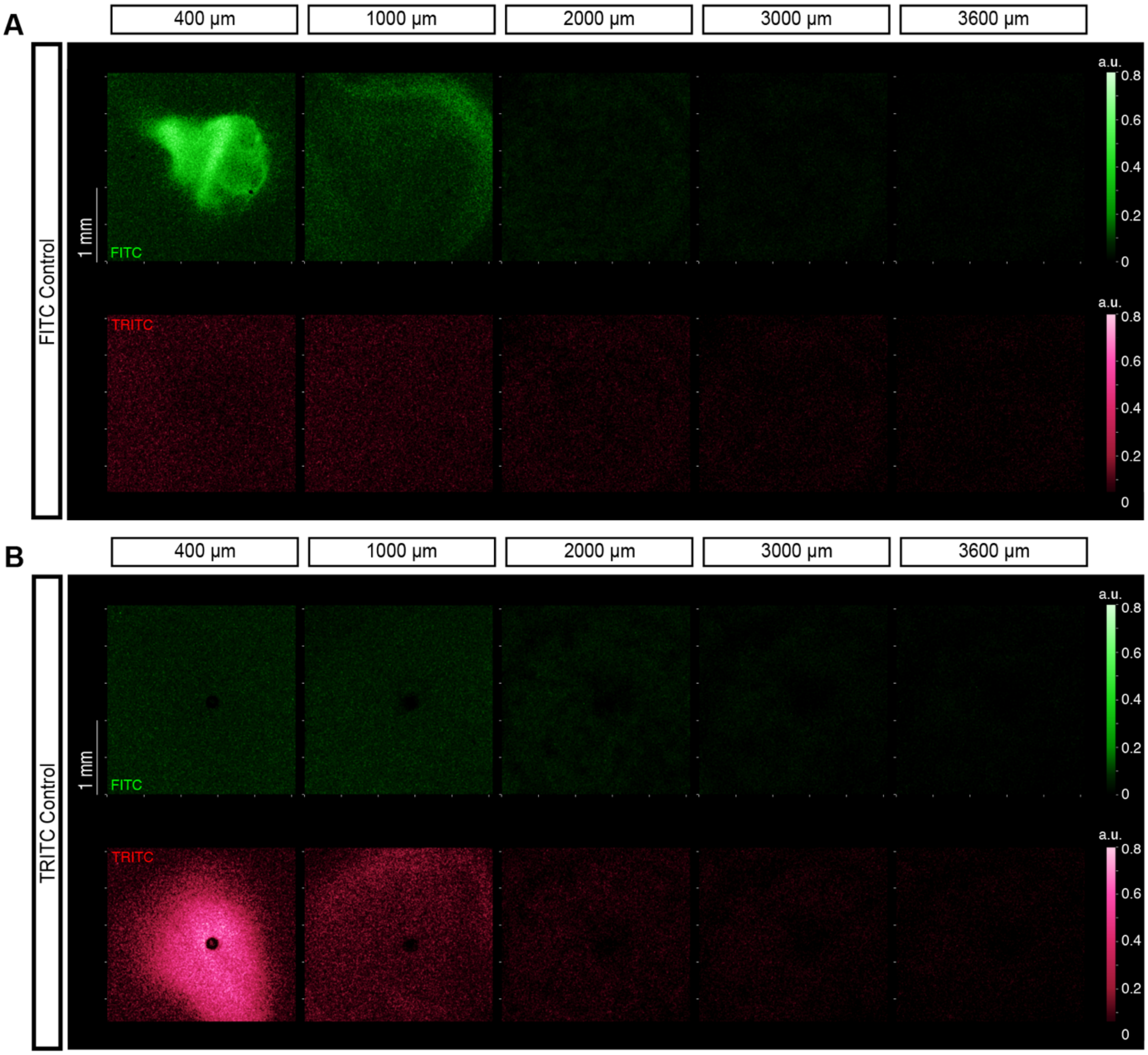


**Figure S17. Representative fluorescence images across different transverse planes for two-morphogen positive controls at Day 0.** Data for (A) FITC-dextran and (B) TRITC-dextran at Z= 400, 1000, 2000, 3000, and 3600 μm are shown. These images were taken from the same hydrogels presented in Figure 6.

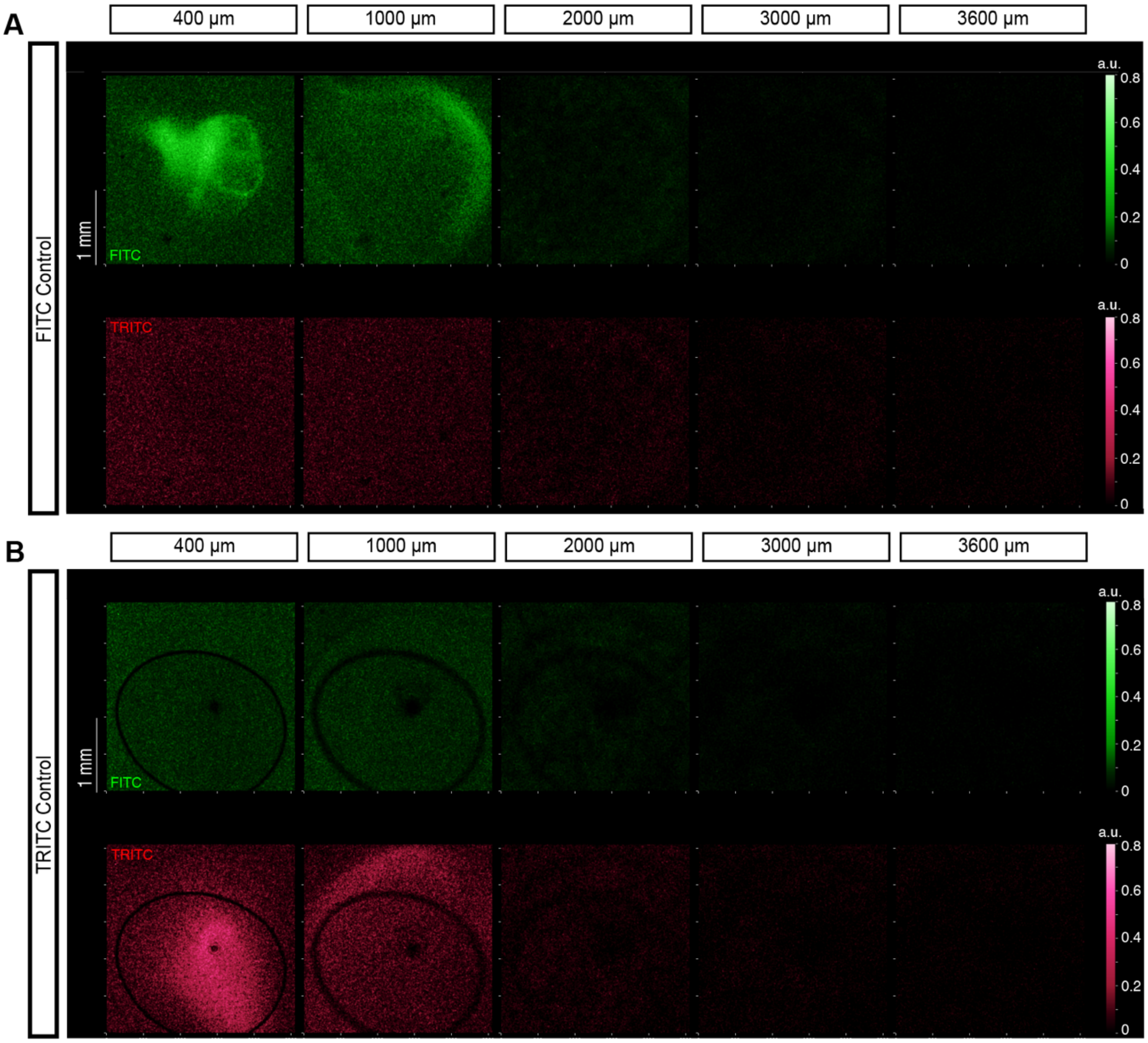


**Figure S18. Representative fluorescence images across different transverse planes for two-morphogen positive controls at Day 1.** Data for (A) FITC-dextran and (B) TRITC-dextran at Z= 400, 1000, 2000, 3000, and 3600 µm are shown. These images were taken from the same hydrogels presented in Figure 6.

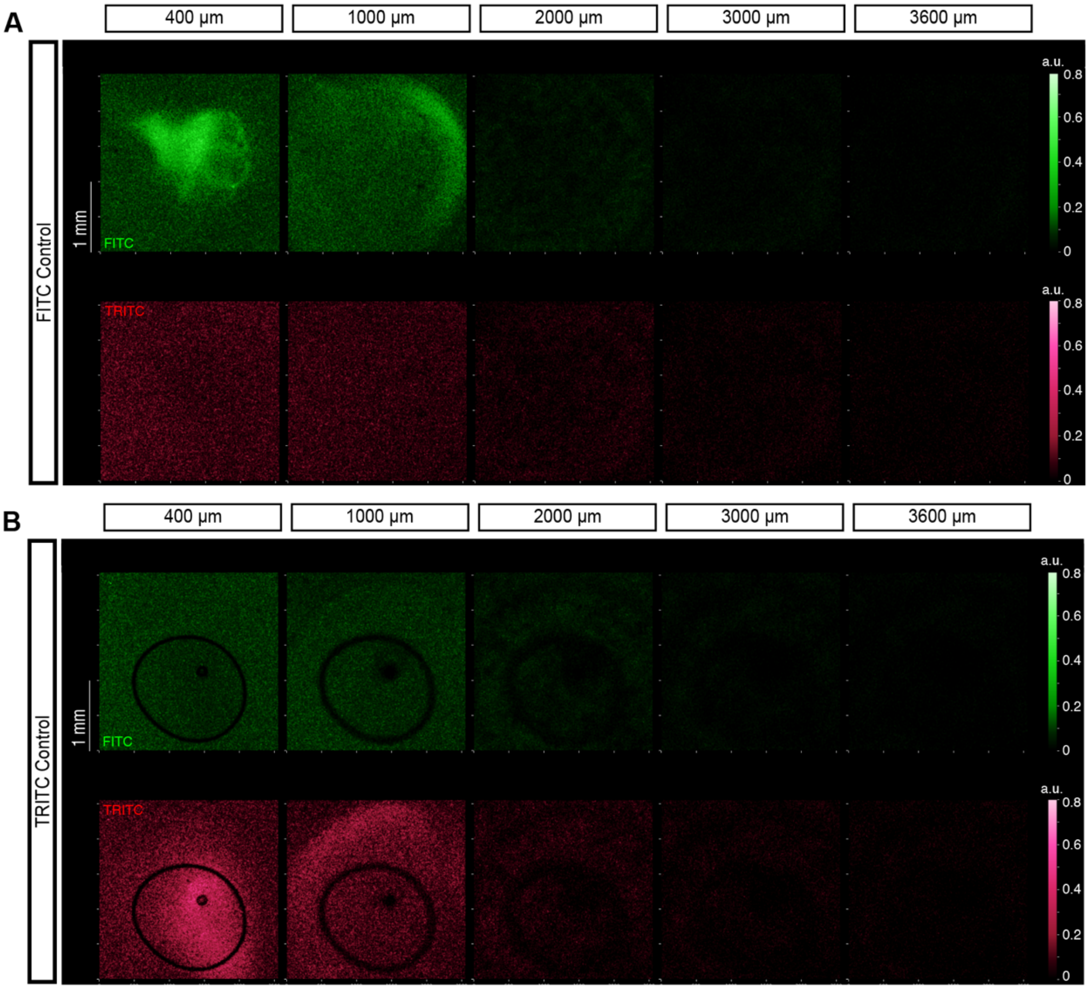


**Figure S19. Representative fluorescence images across different transverse planes for two-morphogen positive controls at Day 3.** Data for (A) FITC-dextran and (B) TRITC-dextran at Z= 400, 1000, 2000, 3000, and 3600 μm are shown. These images were taken from the same hydrogels presented in Figure 6.

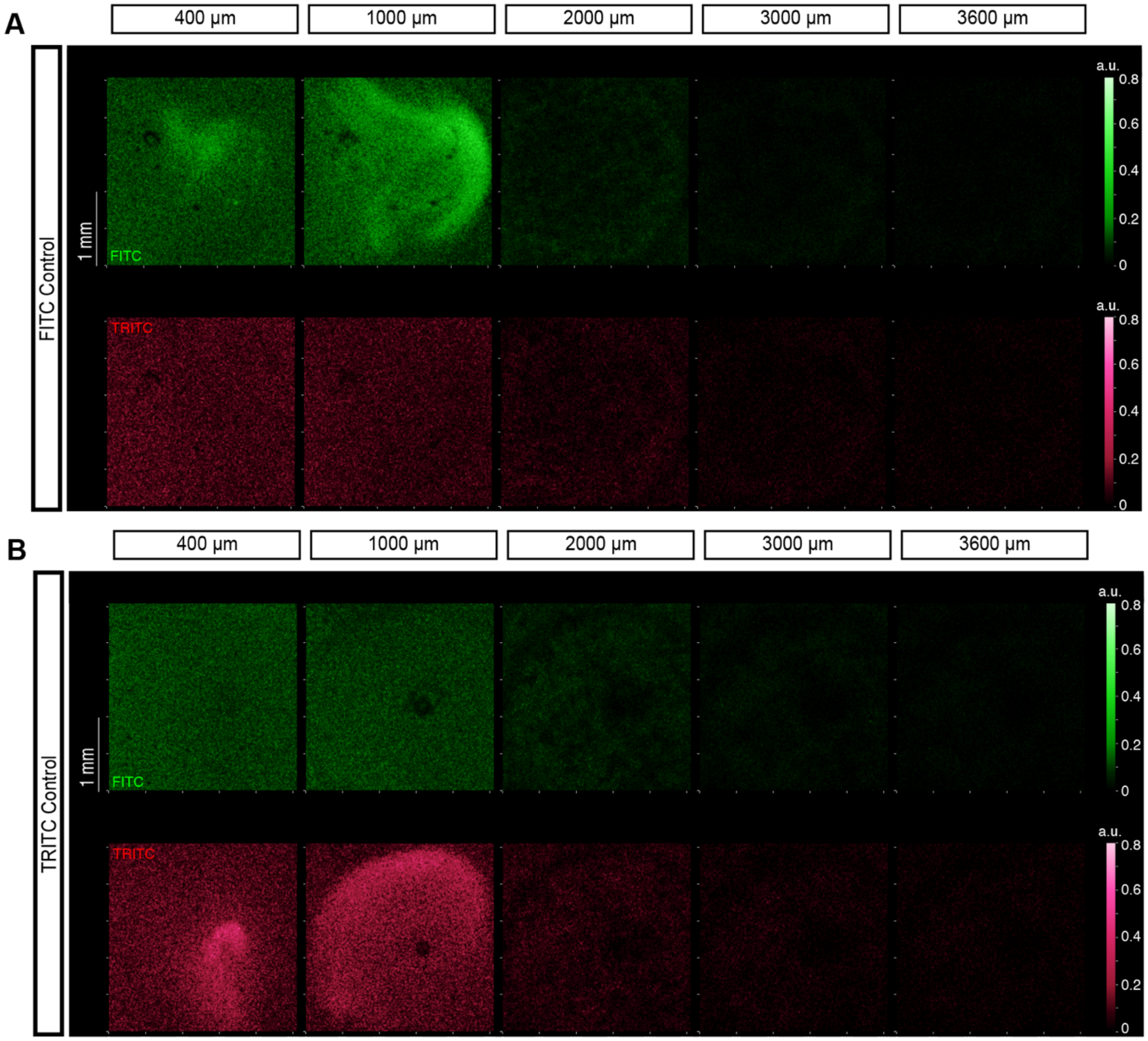


**Figure S20. Representative fluorescence images across different transverse planes for two-morphogen positive controls at Day 6.** Data for (A) FITC-dextran and (B) TRITC-dextran at Z= 400, 1000, 2000, 3000, and 3600 μm are shown. These images were taken from the same hydrogels presented in Figure 6.
